# XIST Self-regulates its Association with THOC2 and the Nuclear Epigenetic Machinery via miR-186 in Alzheimer’s disease

**DOI:** 10.1101/2025.10.05.680468

**Authors:** Somenath Sen, Debashis Mukhopadhyay

**Affiliations:** Biophysical Science Division, Saha Institute of Nuclear Physics, A CI of Homi Bhabha National Institute, Kolkata 700 064, India

**Keywords:** Alzheimer’s disease, XIST, lncRNA, X-chromosome inactivation, miR-186, EZH2, THOC2, ceRNA

## Abstract

Alzheimer’s disease (AD) exhibits a strong female bias, yet the underlying molecular basis remains poorly understood. This study identifies the long non-coding RNA (lncRNA) XIST, a master regulator of X-chromosome inactivation (XCI), as a driver of female-specific AD pathology. Single-nucleus RNA-seq (snRNA-seq) analyses from human AD cortex and *in vitro* AD models reveal elevated and abnormally cytoplasmic XIST, a feature not previously reported in neurodegeneration. Reduced EZH2 levels in AD impair histone H3 lysine-27 trimethylation (H3K27me3) on the inactive X chromosome, disturbing epigenetic silencing. The epigenetic dysregulation elevates the X-linked RNA export factor THOC2, which participates in a feedback loop with XIST. Cytoplasmic XIST functions as a competing endogenous RNA (ceRNA) by sequestering miR-186-5p, thereby rescuing EZH2 and THOC2 transcripts. The resulting XIST/miR-186/EZH2/THOC2 axis couples XIST-driven epigenetic changes with nuclear RNA export pathways via cytoplasmic XIST. Disrupted EZH2-XIST interaction reduces H3K27me3 marks on the inactive X, while increased THOC2 promotes THOC2(TREX)-XIST interaction, maintaining the harmful feedback. Our findings reveal a novel mechanism through which XIST gets recruited to the nuclear RNA export pathways in AD. XIST thus emerges as a critical node in sex-specific AD pathophysiology and a promising target for therapeutic intervention.

**Graphical Abstract:** 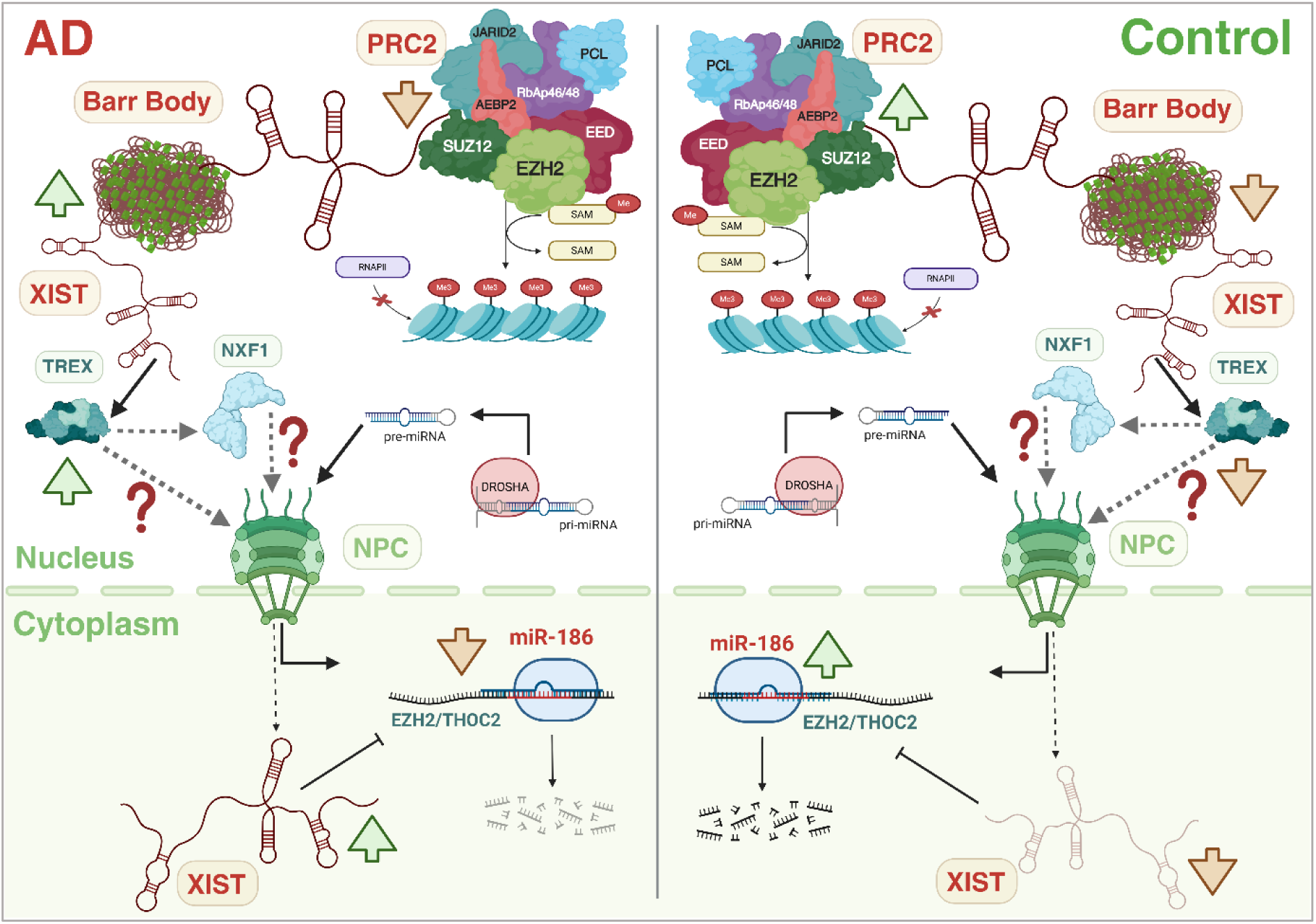

## Introduction

Alzheimer’s disease (AD) is an increasing global health challenge, mainly affecting the elderly and characterized by progressive cognitive decline and memory loss. The disease develops as amyloid β (Aβ) plaques and neurofibrillary tangles (NFTs) accumulate in the brain, leading to neuronal dysfunction and ultimately cell death [1]. Despite extensive research, the exact molecular mechanisms behind AD development are not fully understood, highlighting the need for further investigation into the disease’s underlying regulation.

Recent advances have highlighted the importance of long non-coding RNA (lncRNAs) in influencing various factors behind AD, including gender [2]. Two-thirds of all AD cases are attributed to females [3], and this discrepancy remains unexplained. Recent research has identified sex-specific lncRNA candidates that are dysregulated in AD [2], [4]. One such lncRNA is X-inactive specific transcript (XIST), which plays the major role in X-chromosome dosage compensation [5], is notably upregulated in AD [6]. X-chromosome inactivation is a unique process for maintaining equal expression levels of sex-linked genes, despite females having an extra X chromosome [7]. Consequently, females naturally exhibit higher levels of XIST [8]. Here, we investigate the potential role of XIST as a regulator of the female bias in AD.

XIST is traditionally known to mediate X-inactivation by interacting with the PRC2 complex. A component of this complex, EZH2, functions as a catalytic subunit and promotes the methylation of histone H3 at lysine 27 (H3K27me3) [9]. This histone modification leads to transcriptional repression of the inactivated X chromosome [10]. However, there are more complex roles of XIST beyond X-chromosome inactivation (XCI), especially in the context of various diseases [11], [12], [13]. In proliferative scenarios like many cancer types, XIST acts as an oncogene through its interactions [14], [15], making its upregulation in neurodegeneration particularly unprecedented.

Additionally, challenging the traditional view of XIST’s nuclear localization and retention, a recent study has shown its presence in the cytoplasm under proinflammatory stress [16]. While the cytoplasmic presence of XIST has already been noted in AD [6], the mechanistic details remain unclear. XIST is upregulated and acts as a sponge for miR-186-5p in non-small cell lung cancer, influencing invasion and proliferation [17]. We explored the potential role of the miR-186/XIST axis in AD. Interestingly, in the aging brain, miR-186 is significantly decreased and negatively regulates BACE1 expression, a key gene involved in Aβ production in AD [18]. Although how XIST is present in the cytoplasm is not fully understood, some mRNA export factors are known to facilitate nuclear export of lncRNAs [19]. While reduced NXF1/XIST interaction promotes the nuclear retention of XIST [20], we sought to investigate whether the TREX RNA export complex could participate in overcoming this barrier. THOC2, the largest scaffold subunit of the TREX complex, has been shown to associate with both mRNAs and lncRNAs to facilitate their nuclear export [19], [21]. Notably, the *THOC2* gene itself is an X-linked gene [21], raising the possibility of a regulatory interplay between XIST and the RNA export machinery. Supporting this hypothesis, a previous ChIRP-MS study identified THOC4, another core TREX component, as a XIST-interacting protein [22]. Pull-down assays have demonstrated a direct interaction between THOC4 and THOC2 [23]. Together, these observations suggest that XIST may engage the TREX complex and identify THOC2 as a compelling candidate to investigate whether XIST can non-canonically regulate the expression or function of an X-linked component of the nuclear RNA export machinery.

Here, we demonstrate that the abnormal presence of XIST in the cytoplasm contributes to its overall upregulation in AD and further examine its functional impact on nuclear and cytoplasmic processes. Using an *in vitro* AD model, we show that by sponging miR-186, cytoplasmic XIST preserves EZH2 levels, and reduces THOC2 mRNA degradation, leading to increased THOC2-XIST physical association, which in turn regulates XIST’s own cytoplasmic presence. These novel findings shed light on the sex bias observed in AD.

## Materials and Methods

### Single-nucleus RNA-seq (snRNA-seq) Data Analysis of AD

#### Data acquisition and preprocessing

The raw count matrix and metadata for the human Alzheimer’s disease single-nucleus RNA-sequencing dataset GSE138852 were downloaded from the Gene Expression Omnibus (GEO) [24]. The dataset consisted of eight 10X Genomics libraries (four AD and four control), with each library representing pooled nuclei from two individuals (total n = 16). Analyses were performed using Seurat (v4.0) (https://satijalab.org/seurat/) in R.

Sex information was incorporated into the metadata to generate male and female subsets for downstream analyses. A total of 6,673 AD nuclei and 6,541 control nuclei were analyzed. Female-specific analyses included 2,605 AD nuclei and 1,122 control nuclei.

#### Quality control and preprocessing

Quality control filtering excluded nuclei with fewer than 50 detected genes, more than 5,000 detected genes, or greater than 10% ribosomal transcript content. Nuclei annotated as doublets in the original dataset metadata were excluded prior to downstream analyses. Gene expression values were log-normalized using Seurat’s NormalizeData function, and the 2,000 most variable genes were identified using the variance stabilizing transformation (vst) method.

#### Dimensionality reduction and cell-type annotation

Data were scaled and subjected to principal component analysis (PCA). The first ten principal components were used for nearest-neighbor graph construction, Louvain clustering, and UMAP visualization. Cell identities were assigned using the original dataset annotations together with canonical neuronal markers (RBFOX3/NeuN, ELAVL3 and SYT1). Neuronal nuclei were subsequently extracted for downstream analyses.

#### Differential expression analysis

Differential expression between AD and control neuronal nuclei was performed using Seurat’s Wilcoxon rank-sum test. Benjamini-Hochberg false discovery rate (FDR) correction was applied to all statistical tests, and genes with an adjusted p-value (q-value) < 0.05 were considered statistically significant. Because each sequencing library represented pooled nuclei from two individuals and the female subset contained only one AD and one control pooled library, donor-level pseudo-bulk differential expression analysis, mixed-effect modelling, and covariate-adjusted statistical modelling were not feasible.

#### Functional Enrichment Analysis

Using Pearson’s correlation analysis, the transcriptional changes positively and negatively correlated with XIST expression were calculated. The top 50 correlated genes were visualized using the Pheatmap package in R. Based on these co-expressed genes, the pathways mediated by XIST were enriched through Gene Set Enrichment Analysis (GSEA) with the clusterProfiler package (version 4.14.6) [25] in R. The enriched pathways were then categorized into inhibited and promoted pathways based on their negative (<0) and positive (>0) Normalized Enrichment Score (NES) values, respectively. The top 10 inhibited and promoted pathways were displayed in a dotplot using the ggplot2 package in R [26].

### Female X-chromosome accessibility analysis using single-nucleus ATAC-seq dataset

snATAC-seq data (GSE174367) were analysed using the Signac package in R. Following TF-IDF normalization, feature selection, and latent semantic indexing (LSI), female neuronal nuclei were selected based on the provided cell-type annotations and sample metadata. Chromosome X peaks were identified using Ensembl GRCh38 gene annotations (EnsDb.Hsapiens.v86), and a chromosome X accessibility score was calculated for each cell as the mean TF-IDF-normalized accessibility across all peaks mapping to chromosome X. Promoter regions of X-linked genes were defined as 2 kb upstream and 500 bp downstream of the annotated transcription start site using the promoters() function from the GenomicRanges package. Peaks overlapping these promoter regions were identified using findOverlaps(), and an X-linked promoter accessibility score was calculated for each cell as the mean TF-IDF-normalized accessibility across the promoter-overlapping peaks. Cell-level accessibility scores were compared between AD and control neurons using the Wilcoxon rank-sum test.

### Cell Culture and Plasmid Transfection

The human neuroblastoma cell line SH-SY5Y was obtained from the National Centre for Cell Sciences (NCCS, Pune, India), cell repository. The cells were cultured in Dulbecco’s Modified Eagle Medium/Nutrient Mixture F-12 (DMEM-F12, Gibco) supplemented with 10% fetal bovine serum (FBS, Gibco), following standard protocols provided by the supplier. Cell maintenance was performed under controlled conditions in a humidified incubator (Thermo Scientific) set at 37°C with 5% CO2 atmosphere to ensure optimal growth conditions.

For transfection experiments, cells at 70-80% confluency were chosen to ensure optimal transfection efficiency. Lipofectamine 2000 transfection reagent (Invitrogen) was used for all transfection procedures, with the reagent volume set to twice the amount (in microliters) of the plasmid DNA (in micrograms) as recommended by the manufacturer’s protocol. To evaluate transfection efficiency, green fluorescent protein (GFP)-tagged constructs were introduced, and cellular fluorescence was examined after a 48-hour incubation period using fluorescence microscopy. This quality control step verified successful transfection before moving on to subsequent experiments. All procedures were performed under sterile conditions in a laminar flow hood to maintain cell culture integrity.

### Plasmid Constructs, siRNAs and Antibodies

#### Plasmids

The GFP control plasmid (pGFP-C1, Clontech) and the amyloid precursor protein intracellular domain (AICD) construct cloned into the pGFP-C1 vector (referred to as AICD from here on) were available in the laboratory. These constructs had been previously created and characterized in earlier studies [27], [28], [29], [30]. The GFP plasmid served as the baseline control vector, while the AICD construct contained the APP intracellular domain sequence inserted into the same pGFP-C1 backbone. All the constructs were stored as glycerol stocks at liquid nitrogen. Before transfection, plasmid isolation was performed using the Miniprep plasmid kit (Qiagen, Cat No. # 27104). The integrity of all constructs was confirmed by running the purified plasmids on 0.8% agarose gels.

#### siRNAs

XIST was knocked down using pre-designed siRNAs (Qiagen, Cat No. #1027415). The transfection was performed at a final concentration of 50 nM. THOC2 knockdown was performed using MISSION® esiRNA targeting THOC2 (Sigma Cat No. #EHU145541). For THOC2 knockdown, the esiRNAs were transfected at a final concentration of 40nM. All the transfections were performed using Lipofectamine 2000 (Invitrogen) at 37°C.

#### Antibodies

THOC2 Polyclonal Antibody (Invitrogen, Cat No. #PA5-110485, WB 1:1000, ICC 1:500), Ezh2 (AC22) Mouse mAb (Cell Signaling Technology Cat# 3147, RRID:AB_10694383, WB 1:1000, ICC 1:500), Anti-Histone H3 (tri methyl K27) antibody (Abcam Cat# ab192985, RRID:AB_2650559, WB 1:1000, ICC 1:500), Normal Rabbit IgG (Cell Signaling Technology Cat# 2729, RRID:AB_1031062), Normal Mouse IgG (Cell Signaling Technology Cat# 68860, RRID:AB_3675987), HRP Anti-beta Actin antibody [AC-15] (Abcam Cat# ab49900, RRID:AB_867494, WB 1:7000), Anti-Lamin A + Lamin C antibody [4C11] (Abcam Cat# ab238303, RRID:AB_3722722, WB 1:2000), Anti-GAPDH antibody - Loading Control (Abcam Cat# ab9485, RRID:AB_307275).

### AD Cell Model

The SH-SY5Y human female neuroblastoma cells were transfected with either the GFP control vector or AICD construct using Lipofectamine 2000 transfection reagent according to standard protocols. Synthetic amyloid-β peptide (Aβ_1-42_, Sigma) was prepared by dissolving the lyophilized powder in dimethyl sulfoxide (DMSO) to make a stock solution.

Three hours after transfection, specific treatments were applied: GFP-transfected cells received DMSO, while AICD-expressing cells were treated with 0.5 μM Aβ_1-42_ solution. After a 48-hour incubation under standard culture conditions, all samples were harvested for further experiments. The GFP-transfected, DMSO-treated cells served as the control group, while the AICD-expressing, Aβ_1-42_-treated cells represented the AD cell model, as previously characterized (Supplementary Figure S1) [4], [31], [32]. All experimental conditions were conducted simultaneously with the exact passage numbers and culture conditions to reduce variability.

### RNA Isolation, cDNA Conversion, and Quantitative Real-time PCR

Total RNA was extracted from the harvested cells using TRIzol reagent (Invitrogen, USA) following the manufacturer’s recommended protocol. RNA concentration and purity were measured with a Nanodrop 2000 spectrophotometer (Thermo Scientific), and samples were accepted for further processing only if they showed absorbance ratios ∼ 2.0 at 260/280 and 260/230, indicating minimal protein or organic compound contamination.

For cDNA synthesis, 2 μg of purified total RNA was reverse transcribed for each experimental condition. The reverse transcription process was optimized for different RNA species. The oligo(dT)18 primers (Thermo Scientific) were used for mRNA conversion, random hexamers (Thermo Scientific) were employed for lncRNA templates, and stem-loop primers (Eurofins) were used for miRNA reverse transcription following the pulse PCR methodology [33]. The reactions were conducted using a standardized master mix containing 5× reaction buffer, 10 mM dNTP mix, primers, and reverse transcriptase enzyme (Thermo Scientific).

Quantitative real-time PCR was carried out on a QuantStudio™ 3 Real-Time PCR System (Applied Biosystems) using 2X SYBR™ Green PCR Master Mix (Applied Biosystems) and gene-specific primers (Supplementary Table S1). Appropriate endogenous controls were chosen based on transcript type. The 18S rRNA served as the reference for non-coding RNAs, while β-Actin was used for protein-coding gene normalization. Relative gene expression levels were calculated using the 2^−ΔΔCt^ method, with all reactions performed in technical replicates to ensure consistency. Thermal cycling conditions adhered to manufacturer guidelines, including an initial denaturation at 95°C for 10 minutes, followed by 40 cycles of 95°C for 15 seconds and 60°C for 1 minute, ending with a melt curve analysis to confirm amplification specificity.

### Western Blot

Cells were grown in 6-well plates (Tarsons) and washed twice with 1X PBS before being harvested. The cells were then scraped using a cell scraper and lysed with cell lysis buffer (1M Tris-HCl (pH 7.5), 1N NaCl, 0.5M EDTA, 1M NaF, 1M Na3VO4, 10% SDS, 20mM PMSF, 10% Triton X-100, and 50% glycerol, 1X protease inhibitor), and kept on ice for 30 minutes. After incubation, the lysate was spun at 13,000 RPM at 4°C. The pellet was discarded, and the supernatant was used to measure protein concentration using the Bradford assay. The lysates were then loaded onto an 8% SDS-PAGE gel with gel loading dye containing bromophenol blue. Proteins were transferred from the gel to a PVDF membrane (Millipore, Billerica, MA, USA). The membrane was blocked with 5% non-fat dried milk dissolved in TBST (50 mM Tris-HCl, 150 mM NaCl pH 7.5, with 0.05% Tween 20) for one hour, followed by washing with TBST. It was then incubated overnight at 4°C with primary antibody diluted 1:1000 in TBST with shaking. After overnight incubation, the membrane was washed thrice with TBST and then incubated with secondary antibody in TBST (1:3000) for two hours at room temperature. Following secondary antibody incubation, the membrane was washed thrice with TBST. Bands were visualized using the Super Signal West Pico Chemiluminescent Substrate kit (Thermo Scientific, IL, USA) on an Azure Biosystems Imager.

### Combined Immunocytochemistry & Fluorescence in situ Hybridization of RNA (ICC-FISH)

The ICC-RNA-FISH experiments were conducted following the protocol for adherent cell lines as outlined by Stellaris with modifications. Cells were seeded on a 22 mm coverslip placed in a 35 mm cell culture dish two days before transfections. After transfections and treatments, the cells were first washed twice with autoclaved 1X PBS. The cells were then fixed using 4% paraformaldehyde (Merck) (PFA) for 10 minutes at room temperature. Subsequently, the 4% PFA was discarded, followed by two washes with 1X PBS. Cells were permeabilized with 0.5% Triton X (Promega) for 5 minutes, then washed twice with 1X PBS. This permeabilization step was repeated once more. After the final wash with 1X PBS, the cells were immersed in 500 µl of primary antibody diluted in 1X PBS (1:500). Cells were incubated with the primary antibody for 2 hours, then washed twice with 1X PBS for 5 minutes each. Next, the cells were incubated with secondary antibody diluted in 1X PBS (1:1000) for 2 hours. The secondary antibody solution was discarded, and the cells were washed twice with 1X PBS for 5 minutes each. After the second wash, the cells were immersed in 4% PFA for 10 minutes to stabilize protein-protein interactions. Following this, the cells were washed twice more with 1X PBS.

Next, the cells were submerged in 500 µl of wash buffer A (Stellaris) for 5 minutes. The coverslips were moved to a humidified chamber and placed on top of 100 µl of hybridization solution containing RNA FISH probes diluted in the hybridization buffer (Stellaris) at 1:250. They were then incubated overnight at 37°C. After the overnight incubation, the cells were washed with wash buffer A (Stellaris) for 30 minutes. The coverslips were then mounted on glass slides using 1X PBS containing DAPI (1 µg/µl) as the mounting medium. The coverslips were left to incubate at room temperature for 30 minutes and sealed to prevent drying. The slides were stored at 4°C.

### 3D Colocalization Analysis from Confocal and Super-resolution Imaging Data

ICC-FISH slides were imaged using confocal and super-resolution settings on a Zeiss LSM 880 with Airyscan 2 microscope across multiple z-planes. The raw data from the microscope were analyzed using the Fiji software [34]. First, the ROIs were selected for each colocalizing channel. The selected ROIs were then extracted individually containing the z hyperstacks for each channel. Next, the 3D ROIs were analyzed using the Coloc 2 (https://imagej.net/plugins/coloc-2) plugin in Fiji to calculate the Pearson’s colocalization coefficient (r value). The Z-stack Depth Color Code (https://github.com/UU-cellbiology/ZstackDepthColorCode) plugin was used to determine the depth of each stack for each colocalizing channel.

### Nuclear and Cytoplasmic Fractionation

For each fractionation experiment one 150 mm cell culture dish was used. Cells were scraped with a cell scraper (Tarsons) and washed twice with pre-chilled 1X PBS. Before the final wash, cells were equally distributed between two 1.5 ml microcentrifuge tubes (Tarsons) and marked protein and RNA respectively.

### Protein Extraction

The cell pellets were lysed with freshly prepared ice-cold lysis buffer (20 mM Tris-HCl pH 7.5, 100 mM KCl, 5 mM MgCl₂, 0.3% NP-40) supplemented with protease inhibitor cocktail. The cells were incubated on ice with frequent mixing for 20 minutes. The cells were then passed through a 1 ml syringe with an attached needle to get rid of any remaining unlysed cells. The lysis was confirmed by observing the lysate under an inverted phase-contrast microscope (Zeiss). The nuclei were pelleted by centrifugation by spinning the lysate at 2,500 × g for 10 minutes at 4°C, the supernatant was collected in a separate tube marked as cytoplasm. The nuclear pellet was washed thrice with the ice-cold cell lysis buffer. The nuclei were lysed with 5X Laemmli buffer.

### RNA Extraction

RNA isolation from the nuclear and cytoplasmic fractions of the cells was performed using the RNA Subcellular Isolation Kit (Active Motif) following the manufacturer’s instructions with modifications. All the centrifugation steps were carried out at 4°C in a refrigerated centrifuge (Thermo). Cells were washed twice with pre-chilled 1X PBS and collected using a cell scraper (Tarsons). The detached cells were collected in microcentrifuge tubes and spun at 2,500 RCF for 5 minutes. The supernatant was discarded, followed by a final wash in pre-chilled 1X PBS. The cells in the pellets were then lysed with ice-cold lysis buffer and kept on ice for 20 minutes, mixed by vortexing and inversion every 3 minutes. Proper rupture of the cell membrane and the presence of free nuclei were monitored under an inverted phase-contrast microscope (Zeiss). After incubation, the tubes were spun at 14,000 RPM for 5 minutes to pellet the nuclei. The supernatants containing the cytoplasmic fractions were transferred to a new tube. The nuclear pellet was washed with 200 µl of 70% ethanol at 14,000 RPM for 5 minutes. Next, 250 µl and 350 µl of buffer G containing β-mercaptoethanol (Sigma) for cytoplasmic and nuclear fractions, respectively, were added to their respective tubes. The tubes were vortexed at maximum speed for 30 seconds, then 350 µl of pre-chilled molecular grade 70% ethanol (Sigma) was added to each tube. The solution was homogenized by pipetting up and down using a P200 pipette tip. The nuclear and cytoplasmic lysates were then transferred to their respective purification columns and washed with 650 µl of wash buffer. Following a final wash with pre-chilled 70% ethanol (Sigma), the columns were spun at 14,000 RPM for another 1 minute to remove any residual ethanol. RNA was eluted by adding nuclease-free water (Promega) directly into the columns and spinning at 14,000 RPM for 2 minutes.

### RNA Immunoprecipitation (RIP)

The RIP experiments were carried out following the abcam’s RIP protocol (https://www.abcam.com/protocols/RIP) with some modifications. Cells were harvested by trypsinization and then resuspended in ice-cold PBS, freshly prepared nuclear isolation buffer (1.28 M sucrose, 40 mM Tris-HCl, pH 7.5, 20 mM MgCl₂, 4% Triton X-100), and nuclease free water. The suspension was incubated on ice for 20 minutes with intermittent mixing and vortexing to ensure thorough lysis. The nuclei were then pelleted by centrifugation at 2,500 × g for 15 minutes. The resulting pellet was resuspended in freshly prepared RIP buffer (150 mM KCl, 25 mM Tris, pH 7.4, 5 mM EDTA, 0.5 mM DTT, 0.5% NP-40, 100 U/ml RNase inhibitor, and protease inhibitors) and kept on ice for 20 minutes with frequent mixing and vortexing. Sonication of the nuclear fraction was performed using the following settings: 30% amplitude for 20 seconds with 30-second intervals, repeated three times. The mixture was then nutated at 20 rpm for 90 minutes at 4°C. After nutation, centrifugation was performed at 12,000 × g for 20 minutes at 4°C to obtain the nuclear lysate. The pellet was discarded, and the supernatant was used for protein quantification via the Bradford assay. For each sample, 5 mg of total protein in RIP buffer was combined with 2.5 µg of target antibody or IgG and incubated overnight at 4°C with gentle rotation. Protein A/G magnetic beads (25 µl) (Thermo) were added to each reaction and incubated for 2 hours at 4°C with gentle rotation. Beads were separated on a magnetic rack; the supernatant was discarded. The beads were washed by resuspending them in 500 µl of RIP buffer, with four washes performed in total. Co-precipitated RNAs were extracted by resuspending the beads in 500 µl of TRIzol reagent, following the manufacturer’s protocol.

### Chromatin Immunoprecipitation (ChIP)

ChIP was performed using the SimpleChIP ® Enzymatic Chromatin IP Kit (Cell Signaling Technology, Cat No. #9003) following the manufacturer’ s instructions with modifications. Briefly, cells were crosslinked by adding formaldehyde to a final concentration of 1% and incubated at room temperature for 10 minutes with gentle rocking. Crosslinking was then quenched with 1.25 M glycine (final concentration 0.125 M) for 5 minutes at room temperature. Cells were washed twice with ice- cold 1 X PBS and collected by centrifugation. Cell pellets were lysed with cell lysis buffer supplemented with protease inhibitors and DTT, then incubated on ice for 15 minutes with frequent vortexing to promote lysis. Nuclei were pelleted at 2,500 g for 5 minutes. The nuclear pellet was washed with 1X wash buffer containing DTT. The nuclei were digested with micrococcal nuclease (Takara) at 37 ° C for 20 minutes. After digestion, the pellet was resuspended in 1X ChIP buffer with protease inhibitors. The nuclear pellet was lysed on ice for 15 minutes with frequent vortexing, followed by sonication to shear chromatin to approximately 150-900 bp. Fragmentation was confirmed via agarose gel electrophoresis. The nuclear debris were pelleted at 9,500 g for 10 minutes, and the supernatant was transferred to a new tube. A Bradford assay was used to measure protein concentrations. Equal amounts of chromatin were diluted in 1X ChIP buffer and incubated overnight at 4°C with 2 µg of antibody against H3K27me 3 (Abcam, # ab 192985). Normal rabbit IgG (Cell Signaling Technology) was used in parallel as a negative control. 2% of input chromatin lysate was reserved. Antibody–chromatin complexes were pulled down with protein G magnetic beads by rotating at 4 ° C for 2 hours. Beads were washed sequentially with low-salt and high-salt buffers to remove nonspecific interactions. Bound chromatin and the 2% input samples were eluted in elution buffer using a thermomixer (Labman) for 30 minutes at 1200 RPM at 65°C. The eluted chromatin was digested with proteinase K at 65°C for 2 hours at 1200 RPM to degrade proteins. DNA was purified with spin columns from the kit and eluted in 15 µL of nuclease- free water. Immunoprecipitated DNA was analyzed by quantitative PCR (qPCR) with locus-specific primers and normalized using the 2% input samples.

### Biotinylated miR-186-5p Mimic Overexpression and RNA Pull-down

The 3’ biotinylated miRNA mimic of miR-186-5p was purchased from Qiagen. The mimic was transfected at a final concentration of 60 nM using Lipofectamine 2000 (Invitrogen) transfection reagent according to the manufacturer’s protocol. Cells were harvested 48 hours after transfection. The RNA pull-down was performed following the pre-established protocol [35]. Cells were washed twice with ice-cold 1× PBS and then harvested by trypsinization and scraping, followed by centrifugation at 2,000 × g for 5 minutes at 4 °C. The resulting cell pellets were transferred to pre-chilled microcentrifuge tubes and either processed immediately or stored at −80 °C. 550 µl of freshly prepared lysis buffer (20 mM Tris-HCl pH 7.5, 100 mM KCl, 5 mM MgCl₂, 0.3% NP-40) supplemented with protease and RNase inhibitors was added to each pellet. Cells were lysed by pipetting up and down and incubated on ice for 20 minutes with frequent intermittent mixing to promote lysis. Lysates were spun by centrifugation at 18,000 × g for 10 minutes at 4 °C, and supernatants containing the cytoplasm were collected for subsequent analysis. In parallel, 50 µl of Dynabeads™ MyOne™ Streptavidin T1 beads (Invitrogen) per sample were prepared by sequential washes: twice with Solution A (0.1 M NaOH, 0.05 M NaCl), twice with Solution B (0.1 M NaCl, 70% ethanol), and three times with lysis buffer. Beads were then blocked in 500 µl lysis buffer containing BSA (Merck) and yeast tRNA (Invitrogen) by rotating at 4 °C for 2 hours. After blocking, beads were briefly washed with lysis buffer and incubated with 500 µl of cytoplasmic lysate overnight at 4 °C with gentle rotation. The remaining 50 µl aliquot of each lysate was retained and stored at −80 °C for 10% input RNA extraction. The next day, beads were washed four times with 500 µl of lysis buffer to remove non-specific interactions. For RNA extraction, both the pull-down samples and stored input lysates were mixed with 500 µl TRIzol, then stored at −20 °C till the RNA isolation initiation step.

### Dual-luciferase reporter assay

The wild-type 3′UTR sequences of THOC2 and EZH2, along with their corresponding mutant 3′UTR sequences containing disrupted miR-186 binding sites, were cloned into the psiCHECK-2 dual-luciferase reporter vector (Promega) (Barcode Biosciences). Each insert was ligated into the multiple cloning site immediately downstream of the Renilla luciferase (hRluc) stop codon, thereby serving as the 3′UTR of the Renilla transcript. The resulting constructs were designated psiCHECK2-THOC2-3′UTR, psiCHECK2-THOC2-3′UTR-MUT, psiCHECK2-EZH2-3′UTR, and psiCHECK2-EZH2-3′UTR-MUT.

SH-SY5Y cells were co-transfected with the indicated psiCHECK-2 reporter constructs together with either a miR-186 mimic or the corresponding negative control using Lipofectamine 2000 according to the manufacturer’s instructions. Forty-eight hours after transfection, cells were lysed in 1X Passive Lysis Buffer, and Firefly and Renilla luciferase activities were measured sequentially using the Dual-Luciferase® Reporter Assay System (Promega) according to the manufacturer’s instructions. Renilla luciferase activity was normalized to Firefly luciferase activity Renilla/Firefly ratio. The mean Renilla/Firefly ratio of the corresponding miR-negative control group was used as the normalization factor. Individual Renilla/Firefly values for both the WT and MUT reporter constructs in the miR-186-treated groups were divided by the corresponding miR-NC group mean prior to statistical analysis.

### Statistical Analysis

Statistical analyses were conducted to assess the significance of the experimental results. For comparisons between two groups, an unpaired student’s t-test and in case of more than two groups one-way ANOVA followed by Bonferroni post hoc test was performed using GraphPad Prism (version 9), available at (https://www.graphpad.com/). The p-values from snRNA-seq data analyses were generated by the Seurat package (v4) (https://satijalab.org/seurat/). All statistical tests were carried out with at least three independent biological replicates to ensure the reliability of the results.

The threshold for statistical significance was set as follows: results with p-values of 0.05 or higher were considered statistically non-significant and labeled as “ns” (not significant). Results with p-values less than 0.05 were marked with a single asterisk (*), indicating significance at the 95% confidence level. P-values below 0.01 were denoted by double asterisks (**), those under 0.001 were shown with triple asterisks (***), and those under 0.0001 were shown with quadruple asterisks (****). This notation system was applied consistently across all figures and tables to clearly indicate the statistical significance of all reported results.

## Results

### XIST upregulation in the AD brain regulates synaptic transmission and transcriptional repression

The snRNA-seq analysis pipeline was used to examine XIST expression across different cell types in female AD and control brain samples from the GSE138852 dataset. Neuronal nuclei were confirmed through enrichment of canonical neuronal markers and the dataset metadata. Visualization on the UMAP plot showed colocalization of XIST with ELAVL3, demonstrating XIST expression in neuron-derived nuclei (Figure 1.A).

**Figure 1.**
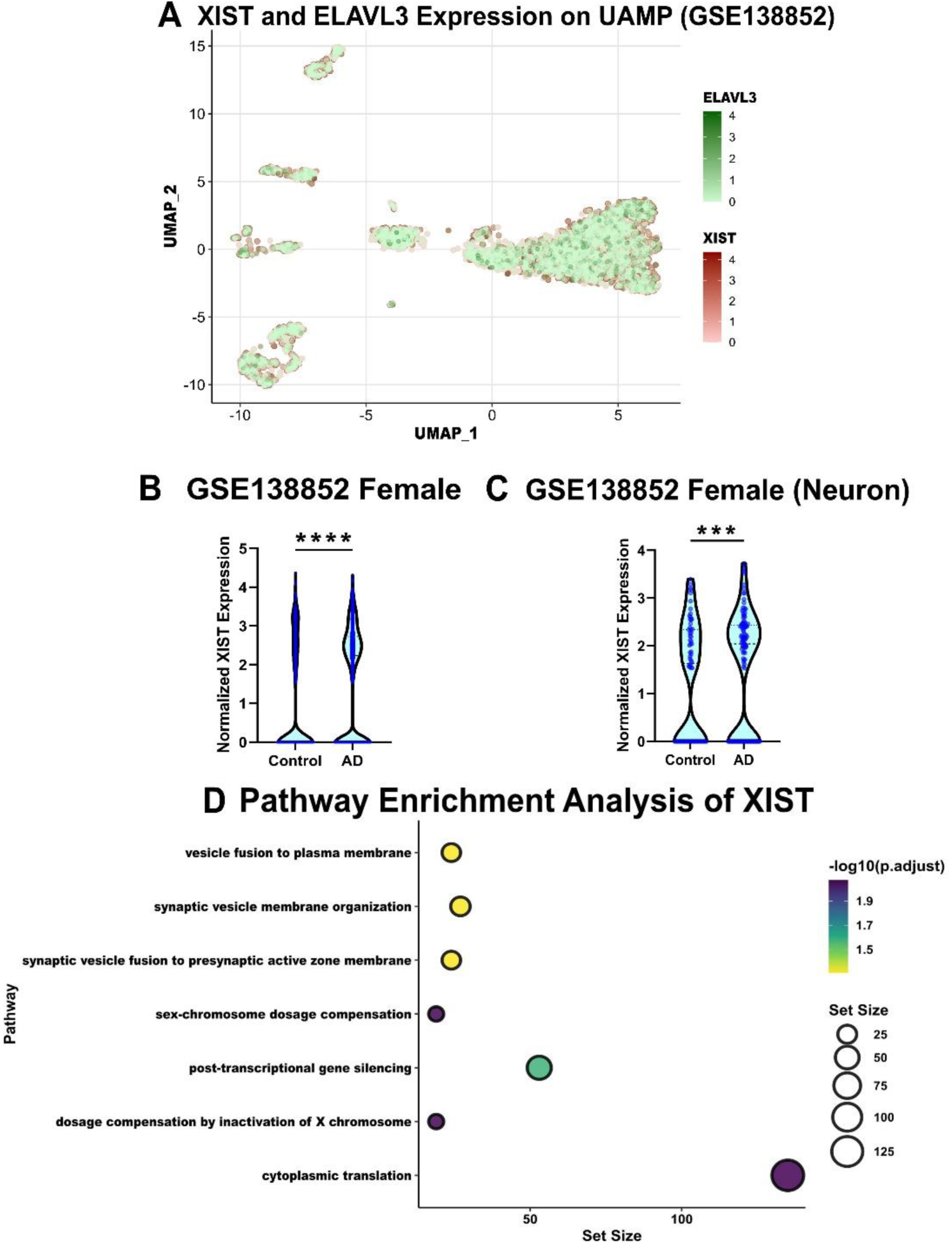
(A) UMAP projection of female nuclei from the GSE138852 snRNA-seq dataset showing the expression of XIST overlaid with the neuronal marker ELAVL3, demonstrating neuronal XIST-expressing cells. (B) Violin plot showing significantly increased normalized XIST expression in female AD nuclei compared with female control nuclei. (C) Violin plot showing significantly increased normalized XIST expression specifically in female neuronal nuclei from AD compared with controls. (D) Gene Set Enrichment Analysis (GSEA) of XIST-associated genes in female AD neurons, highlighting enrichment of pathways related to cytoplasmic translation, post-transcriptional gene silencing, X-chromosome dosage compensation, and synaptic vesicle organization and fusion.

The female neuronal subset demonstrated detectable XIST expression in neuronal nuclei, with higher cell-level XIST expression observed in AD compared with control nuclei in the GSE138852 dataset (Figure 1.B-C).

To evaluate the biological implications of XIST upregulation in AD, Pearson’s correlation analysis was conducted, revealing genes that are co-expressed with increased XIST levels in female neurons (Supplementary Figure S2.).

Subsequent Gene Set Enrichment Analysis (GSEA) of the co-expressed genes revealed biological pathways linked to changes in XIST expression. Pathways with significant negative enrichment scores included cytoplasmic translation (GO:0002181), synaptic vesicle fusion with the presynaptic active zone membrane (GO:0031629), and synaptic vesicle membrane organization (GO:0048499), indicating a decrease in synaptic and translational activities. Conversely, pathways associated with high XIST expression that showed positive enrichment scores included, sex-chromosome dosage compensation (GO:0007549), and post-transcriptional gene silencing (GO:0016441), suggesting increased heterochromatinization and RNA-mediated silencing (Figure 1.D).

Taken together, these analyses suggest that XIST upregulation in AD neurons may contribute to impaired synaptic function and decreased protein synthesis, while also promoting transcriptional repression and epigenetic remodeling. This positions XIST as a potential regulator of neuronal dysfunction in AD.

### XIST shows aberrant nucleocytoplasmic upregulation in AD

To gain a better understanding of its role, XIST expression and its subcellular localization were examined in an *in vitro* AD model (Figure 2.A) using subcellular fractionation followed by qPCR and western blots and RNA-FISH studies.

**Figure 2.**
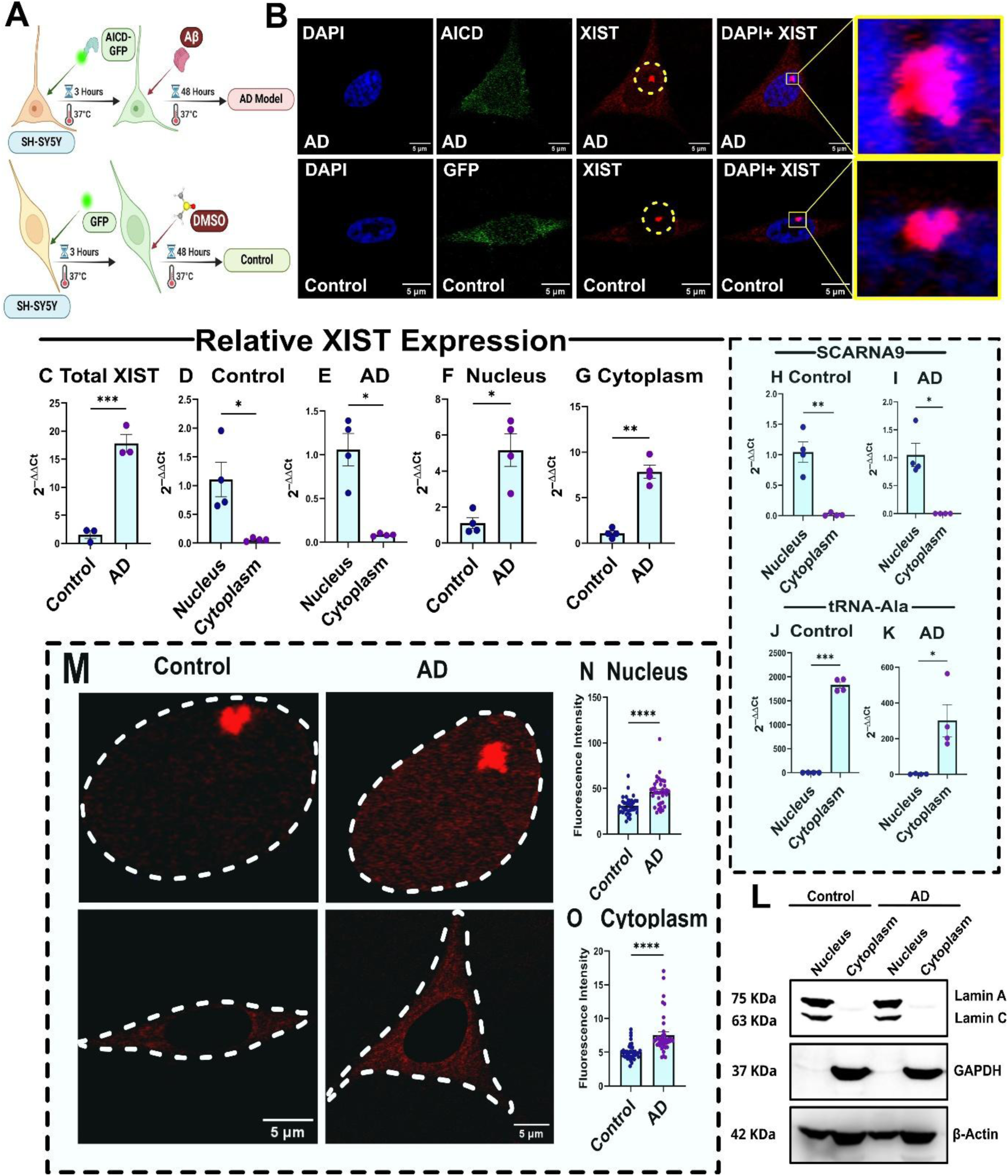
XIST ectopic distribution in the AD cell model. (A) Schematic of the experimental workflow. SH-SY5Y female neuroblastoma cells were transfected with AICD-GFP or GFP constructs and incubated for 3 h at 37°C, followed by treatment with Aβ peptide (AD model) or DMSO (control) for 48 h. (B) Representative confocal images showing subcellular localization of XIST RNA (red puncta) in AD and control cells. DAPI (blue) stains nuclei, while AICD-GFP or GFP (green) indicates successful transfection. Right panels show magnified views of the XIST signal. (C) Total XIST expression was significantly increased in AD cells compared with controls. (D, E) Relative nuclear and cytoplasmic XIST expression in control and AD cells, respectively. (F, G) XIST levels in the nuclear and cytoplasmic fractions were significantly increased in AD compared with controls. (H, I) RT-qPCR analysis of the nuclear-retained RNA SCARNA9 in control and AD cells confirmed efficient subcellular fractionation. (J, K) RT-qPCR analysis of cytoplasmic tRNA-Ala in control and AD cells further validated fractionation efficiency. (L) Western blot validation of nuclear and cytoplasmic fractions using Lamin A/C as the nuclear marker and GAPDH as cytoplasmic markers. (M) Representative RNA-FISH images showing XIST localization in control and AD cells. Dashed white lines outline the cellular and nuclear boundaries. (N, O) Quantification of XIST fluorescence intensity in the nuclear and cytoplasmic compartments revealed significantly elevated XIST signals in both compartments in AD cells. RT-qPCR data are presented as 2^−ΔΔCt and fluorescence intensity data are presented as mean ± SEM. *p < 0.05, **p < 0.01, ***p <0.001, ****p <0.0001.

With RNA-FISH followed by confocal imaging, it was observed that the most intense XIST signals were concentrated on the inactivated X chromosome, appearing as a distinct bright red punctum in the nucleus that co-localized with DAPI. Interestingly, a notable presence of XIST in the cytoplasm was also detected (Figure 2.B); an atypical and previously uncharacterized localization in the context of AD. This unusual cytoplasmic presence suggests a potentially novel function of XIST in the disease pathology.

The qPCR analysis confirmed that total XIST expression was significantly upregulated in the AD model (Figure 2.C). To further clarify this upregulation, subcellular fractionation combined with RNA-FISH-based intensity measurements were used to separately quantify nuclear and cytoplasmic XIST levels. In both control and AD cells, nuclear XIST levels were substantially higher than cytoplasmic levels. However, when comparing nuclear and cytoplasmic XIST levels between AD and control conditions, it was observed that, along with the significant increase in nuclear XIST in AD, cytoplasmic XIST levels were also markedly elevated compared to controls. Interestingly, the upregulation in the cytoplasmic XIST was more compared to the nuclear transcript pool (Figure 2.D-G). The SCARNA9 RNA expression was mainly localized to the nucleus, while tRNA expression was predominantly found in the cytoplasm, indicating effective fractionation (Figure 2.H-K). Lamin A/C was used to further validate the fractionation efficiency of the nuclear fractions, while GAPDH served as the cytoplasmic marker, both further confirming the purity of the fractionated samples (Figure 2.L).

To further examine XIST distribution, confocal images from RNA-FISH experiments were analyzed by segmenting individual cells based on DAPI signals, differentiating nuclear from cytoplasmic regions. Consistent with our qPCR results, image-based intensity measurements showed that both the nuclear and cytoplasmic XIST signal was significantly higher in AD compared to control (Figure 2.M-O). These findings collectively demonstrate that XIST is upregulated in both the nuclear and cytoplasmic compartments in AD. Although nuclear XIST remains the predominant transcript pool, the cytoplasmic fraction exhibits a proportionally greater increase relative to control, suggesting that cytoplasmic XIST may represent novel component of XIST dysregulation in AD.

### EZH2-XIST association is altered in AD, leading to reduced H3K27me3 marks on the inactivated X chromosome

The altered nucleocytoplasmic levels of XIST observed in AD led to two different aspects of its function; its traditional nuclear role and a new cytoplasmic localization. We started by focusing on nuclear XIST to examine its importance in the AD context.

In our AD model, we measured EZH2 expression at both mRNA and protein levels and found it to be significantly decreased (Figure 3.A-C). Therefore, we wanted to determine if the quantity of XIST bound to EZH2(PRC2) is also altered. To do this, we performed RIP-qPCR with an anti-EZH2 antibody. The pull-down experiments confirmed a significant reduction in EZH2 bound XIST in AD conditions (Figure 3.D-F). This spatial relationship was further validated using combined immunocytochemistry (ICC) and RNA-FISH with an anti-EZH2 antibody and ssDNA FISH probes targeting XIST (Figure 4.A-B). The 3D imaging analysis showed a noticeably decreased colocalization between XIST and EZH2 in AD cells compared to controls (Figure 4.E).

**Figure 3.**
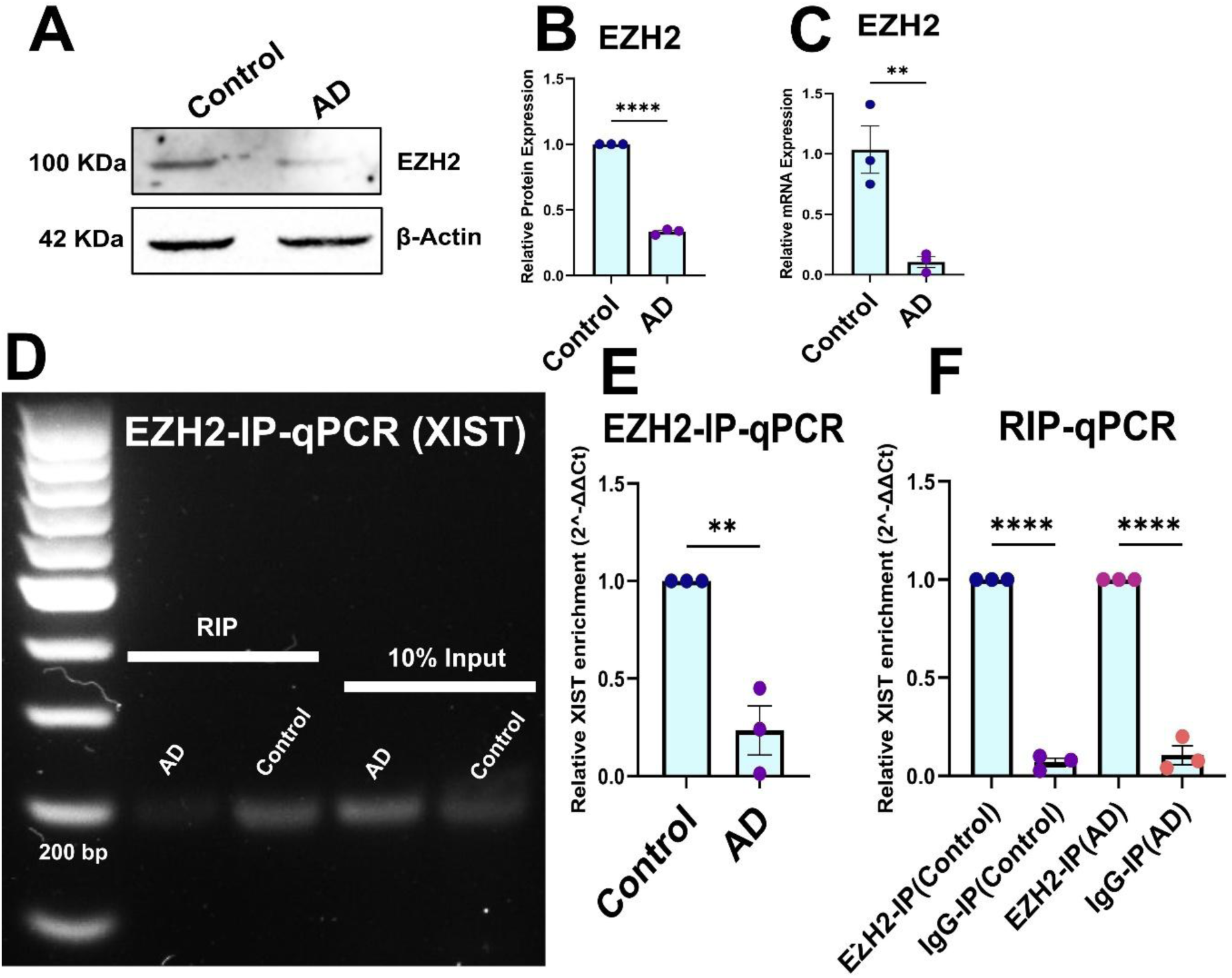
Impaired EZH2 expression and reduced EZH2-bound XIST in AD. (A) Representative immunoblot showing reduced EZH2 protein expression in AD-treated cells compared with controls. β-Actin served as the loading control. (B) Quantification of EZH2 protein levels revealed a significant decrease in AD-treated cells. (C) qRT-PCR analysis indicated a significant reduction in EZH2 mRNA expression in AD-treated cells relative to controls. (D) Representative agarose gel of EZH2-RIP-qPCR showing reduced XIST in EZH2-immunoprecipitated RNA in AD cells compared to control. (E) RIP-qPCR confirmed a significant decrease in XIST RNA bound to EZH2 in AD-treated cells compared with controls. (F) Enrichment of XIST RNA in EZH2-IP compared with IgG-IP in both GFP+DMSO-treated control cells and AICD+Aβ-treated AD cells validated the specificity of the RIP assay and confirmed the interaction between XIST and EZH2 under both conditions. Data are presented as mean ± SEM.**p < 0.01, ****p <0.0001 (unpaired t-test).

**Figure 4.**
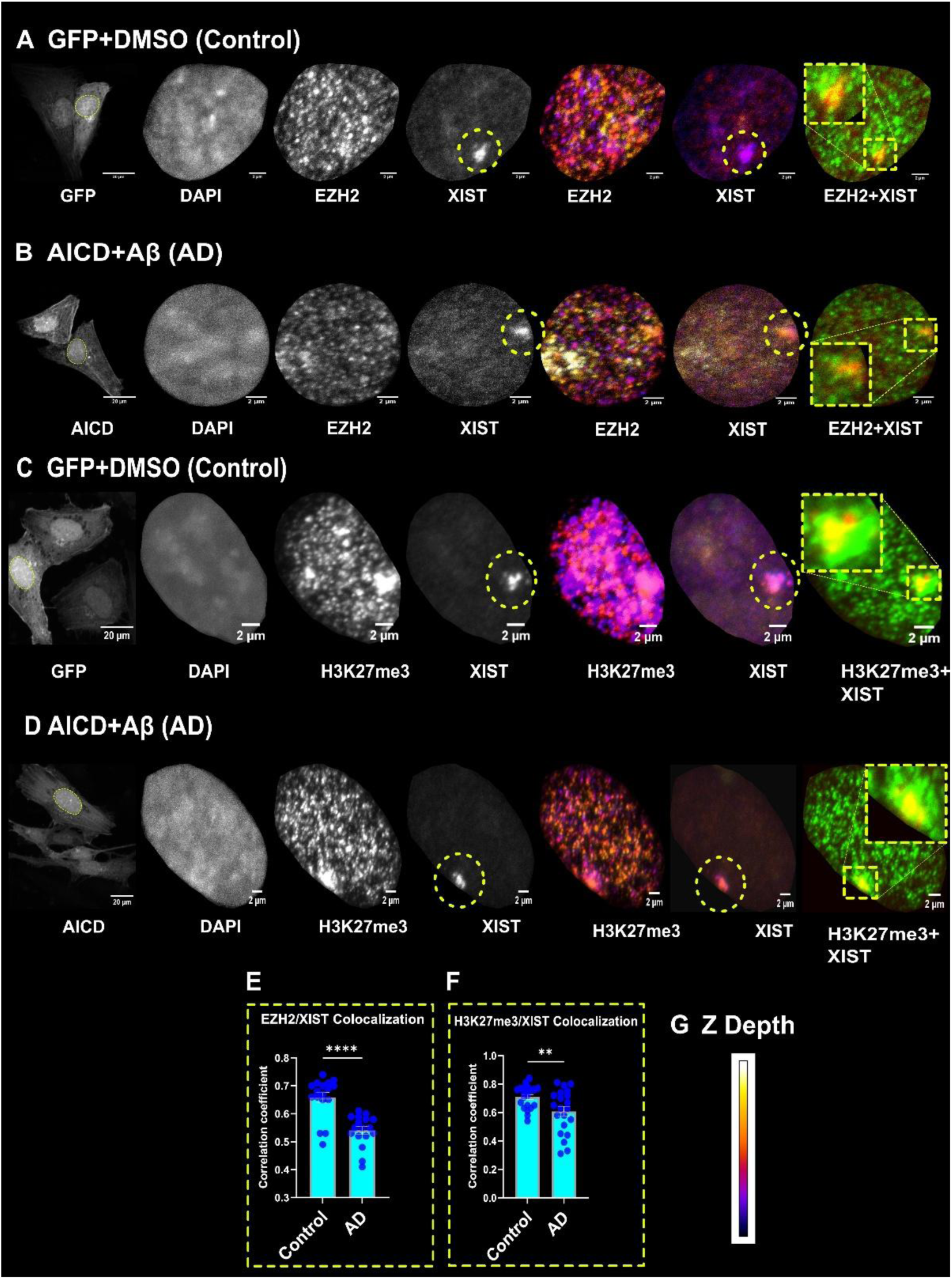
Reduced colocalization of XIST with EZH2 and H3K27me3 in the AD model. (A) Representative confocal images of GFP+DMSO-treated control cells showing nuclear localization of EZH2 and XIST. The merged EZH2+XIST image demonstrates strong colocalization between the two signals. (B) Representative confocal images of AICD+Aβ-treated AD cells showing reduced colocalization between EZH2 and XIST. (C) Representative confocal images of GFP+DMSO-treated control cells showing nuclear localization and colocalization of XIST with H3K27me3, a repressive histone modification catalyzed by EZH2. The merged H3K27me3+XIST image demonstrates a high degree of spatial overlap. (D) Representative confocal images of AICD+Aβ-treated AD cells showing reduced colocalization between XIST and H3K27me3 despite both signals remaining detectable within the nucleus. Magnified insets highlight regions of signal overlap. DAPI staining was used to visualize nuclei. Z-depth color coding was used to depict the axial distribution of fluorescence signals, enabling three-dimensional assessment of colocalization. (E) Pearson’s correlation coefficient analysis demonstrating a significant reduction in EZH2–XIST colocalization in AD cells compared with controls, indicating impaired association of XIST with EZH2 in the AD model. (F) Pearson’s correlation coefficient analysis showing significantly decreased colocalization between XIST and H3K27me3 in AD cells, indicating impaired association of XIST with the repressive H3K27me3 chromatin mark. (G) Z-depth scale used for three-dimensional visualization of fluorescence signal distribution. Error bars represent mean ± SEM.**p < 0.01, ****p <0.0001 (unpaired t-test).

Given the reduced EZH2 bound XIST, subsequent investigation focused on the downstream heterochromatin dynamics mediated by H3K27me3. The overall levels of H3K27me3 were significantly decreased in AD (Supplementary Figure S3). Next, three-dimensional colocalization analysis was performed on z-stacks from dual-labeling experiments combining H3K27me3 immunostaining with XIST RNA-FISH (Figure 4.C-D). This spatial analysis revealed a substantial disruption of the XIST-H3K27me3 association in the AD model. In control cells, XIST RNA foci strongly colocalized with regions enriched for H3K27me3 within the nucleus, as shown in merged confocal z-projections. These colocalized domains, indicated by the overlap of XIST and H3K27me3 signals, demonstrated proper colocalization of XIST to the repressive chromatin compartments of the inactive X chromosome, consistent with its canonical role in gene silencing (Figure 4.C). In AD-mimicking cells in contrast, although H3K27me3 puncta and nuclear XIST signal overlap remained visible (Figure 4.D)., quantitative analysis using Pearson’s correlation coefficient demonstrated a statistically significant decrease in XIST-H3K27me3 colocalization in AD cells compared to controls (Figure 4.F).

The reduction of EZH2 bound XIST not only signals impaired recruitment of PRC2 machinery but also could explain the reduced H3K27me3 colocalization on the barr body.

### Female AD neurons exhibit increased chromosome X chromatin accessibility

Given the reduced association of XIST with EZH2 and H3K27me3 in AD, we next investigated whether these changes were accompanied by altered chromatin accessibility of the inactive X chromosome. To this end, we analyzed a published female single-nucleus ATAC-seq dataset (GSE174367) and quantified chromosome X accessibility specifically in neuronal nuclei. UMAP visualization demonstrated a widespread increase in chromosome X accessibility across female AD neurons compared with control neurons (Figure 5.A). Quantitative analysis further showed that the mean chromosome X accessibility score was significantly elevated in AD neurons (Figure 5.B). Since promoter accessibility is closely associated with transcriptional regulation, we next quantified accessibility across promoter regions of X-linked genes. Consistent with the global chromosome X accessibility analysis, X-linked promoter accessibility was also significantly increased in AD neurons compared with controls (Figure 5.C).

**Figure 5.**
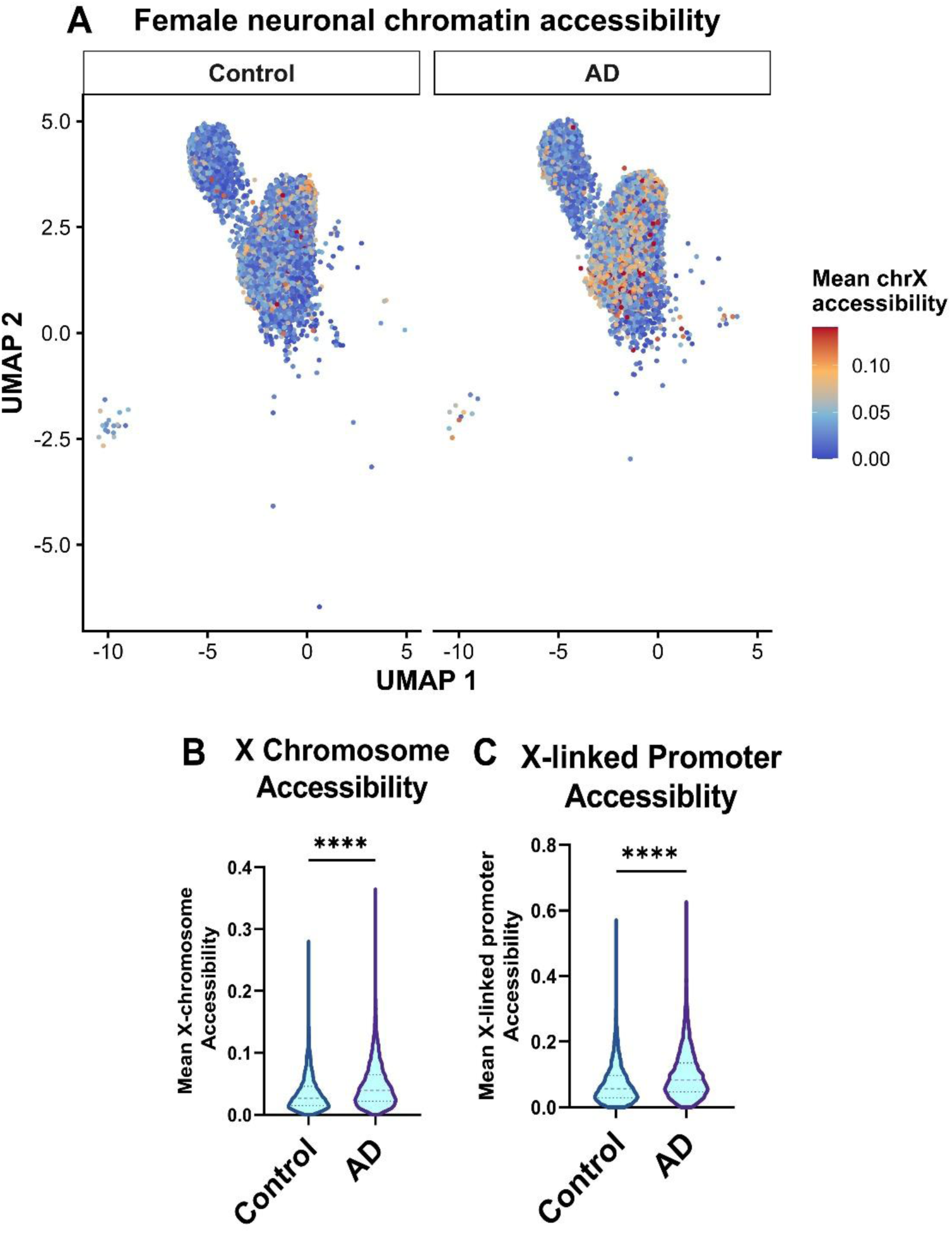
Increased X chromosome chromatin accessibility in female AD neurons. (A) UMAP visualization of female neuronal nuclei from the GSE174367 single-nucleus ATAC-seq dataset showing chromosome X accessibility in control and AD samples. Cells are colored according to the mean chromosome X accessibility score. (B) Violin plot showing significantly increased mean chromosome X accessibility in female AD neurons compared with controls. (C) Violin plot demonstrating significantly increased accessibility at promoter regions of X-linked genes in female AD neurons relative to controls. These findings indicate enhanced chromatin accessibility across the X chromosome, including regulatory promoter regions, in female AD neurons. Data are presented as mean ± SEM. ****p < 0.0001 (Wilcoxon rank-sum test).

These findings indicate that female AD neurons exhibit enhanced chromatin accessibility across the X chromosome, including regulatory promoter regions, consistent with the reduced XIST/H3K27me3 colocalization observed in our experimental model (Figure 4.F). However, because the snATAC-seq dataset does not distinguish the active and inactive X chromosomes, the extent to which the observed increase in accessibility originates specifically from the inactive X chromosome (Xi) cannot be determined.

### Heightened THOC2 levels facilitate THOC2(TREX)-XIST association in AD

THOC2, a protein subunit of the TREX complex and an RNA export factor, has been hypothesized to play a role in the nuclear export of lncRNAs [19]. *THOC2* being X-linked [21], we questioned whether the XIST-mediated altered nuclear epigenetic machinery could influence *THOC2* promoter enrichment in AD through H3K27me3 marks.

To investigate any changes in the chromatin landscape of the *THOC2* promoter region, which has been shown to be silenced in the human Barr body [36], ChIP-qPCR was performed. The results showed that the *THOC2* proximal promoter region has a significantly higher association with repressive H3K27me3 marks in AD compared to the controls (Figure 6.A-C). While the THOC2 mRNA levels were found to be significantly downregulated, interestingly, the THOC2 protein levels were found to be significantly upregulated in AD (Figure 6.D-F).

**Figure 6.**
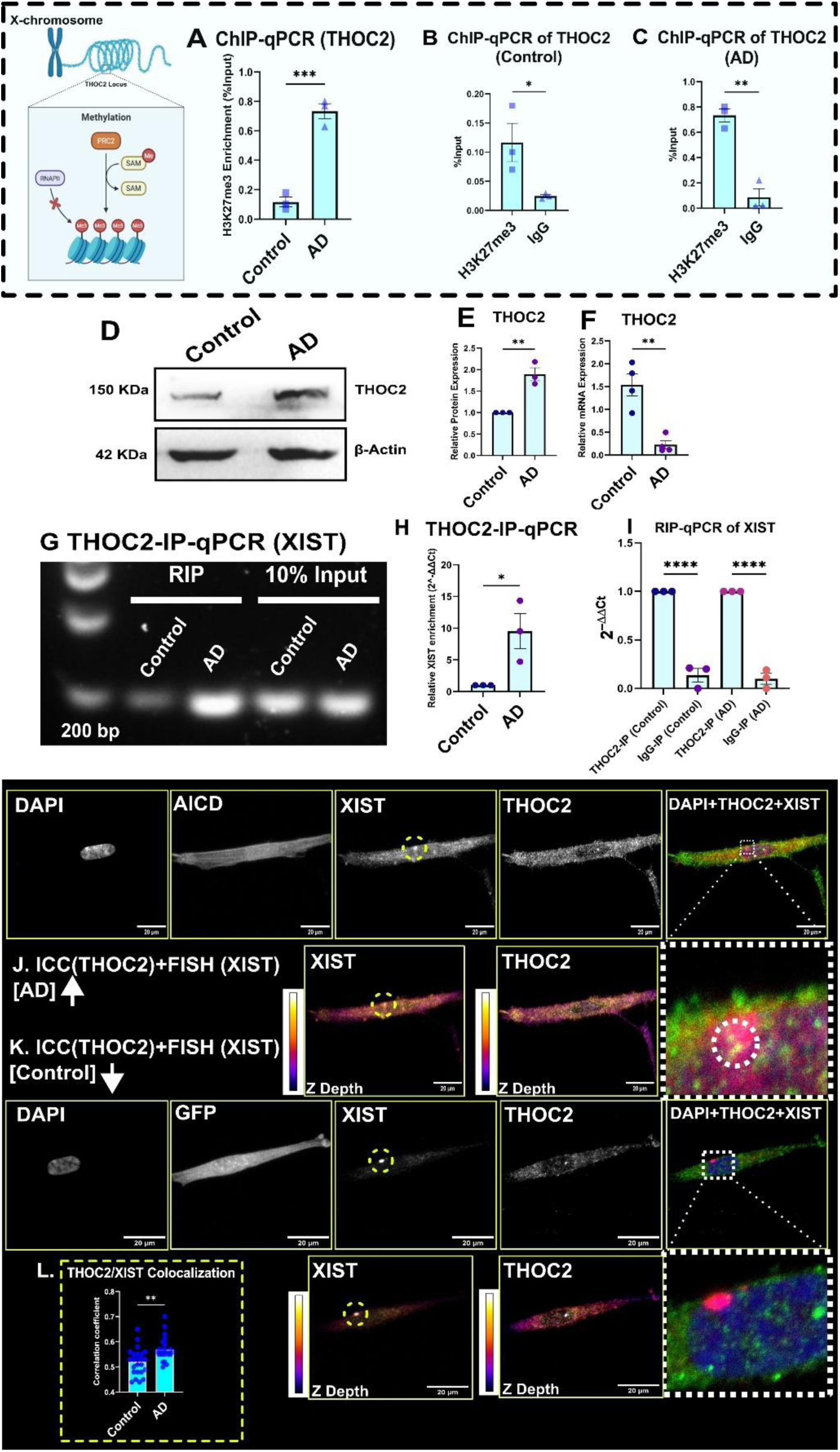
Upregulation of THOC2 and enhanced XIST–THOC2 association in AD. (A) ChIP-qPCR analysis showing increased H3K27me3 enrichment at the THOC2 promoter in AICD+Aβ-treated AD cells compared with GFP+DMSO-treated controls. (B, C) ChIP-qPCR demonstrating specific enrichment of H3K27me3 over IgG at the THOC2 promoter in control and AD cells, respectively. (D) Representative immunoblot showing THOC2 protein expression in control and AD-treated cells. β-Actin served as the loading control. (E,F) Quantification of THOC2 protein expression revealed significant upregulation in AD-treated cells, whereas qRT-PCR analysis demonstrated significantly reduced THOC2 mRNA expression in AD relative to controls. (G) Representative agarose gel of THOC2-RIP-qPCR showing XIST amplification in THOC2-immunoprecipitated RNA AD cells compared to control. (H) RIP-qPCR demonstrating significantly increased association of XIST with THOC2 in AD-treated cells compared with controls. (I) Enrichment of XIST RNA in THOC2-IP compared with IgG-IP in both GFP+DMSO-treated control cells and AICD+Aβ-treated AD cells confirmed the specificity of the RIP assay. (J) Representative confocal images of AICD+Aβ-treated cells showing immunocytochemical detection of THOC2 and RNA-FISH detection of XIST. DAPI staining was used to visualize nuclei. Merged images demonstrate increased colocalization of THOC2 and XIST in AD cells, with enlarged insets highlighting regions of signal overlap. (K) Representative confocal images of GFP+DMSO-treated control cells showing comparatively weaker colocalization between THOC2 and XIST. Z-depth color coding was used to depict the axial distribution of fluorescence signals for three-dimensional assessment of colocalization. (L) Pearson’s correlation coefficient analysis demonstrating significantly increased colocalization between THOC2 and XIST in AD cells compared with controls, indicating enhanced association of XIST with the TREX component THOC2 in the AD model. Data are presented as mean ± SEM. *p < 0.05, **p < 0.01, ***p <0.001, ****p <0.0001 (unpaired t-test).

Next, we wanted to investigate whether this aberrant THOC2 upregulation could lead to any direct interaction with XIST. The RIP-qPCR studies showed increased THOC2-bound XIST in AD compared to control (Figure 6.G-I). These results indicate that the increased abundance of THOC2 protein in AD is accompanied by increased recovery of THOC2-bound XIST in the RIP assay. This suggests that greater availability of THOC2 protein may facilitate the recruitment of XIST to the TREX-mediated RNA export machinery.

Next, using 3D colocalization analysis, we examined the interaction between THOC2 and XIST based on ICC-FISH imaging data in both control and AD cell models (Figure 6.J-K), and we observed a higher colocalization of THOC2-XIST signals in AD compared to controls (6.L).

### THOC2 regulates the nucleocytoplasmic distribution of XIST in AD

The heightened THOC2-bound XIST levels suggested that the TREX-mediated RNA export machinery could be involved in regulating XIST’s cytoplasmic localization. To investigate this, THOC2 was knocked down in the AD condition, followed by fractionation of the cells (Figure 7A-D, Supplementary Figure S4). Subcellular fractionation followed by RT-qPCR demonstrated that THOC2 depletion caused a significant reduction in cytoplasmic XIST accompanied by its subsequently increased nuclear pool, whereas AD cells transfected with the negative control exhibited the opposite distribution (Figure 7E-H).

**Figure 7.**
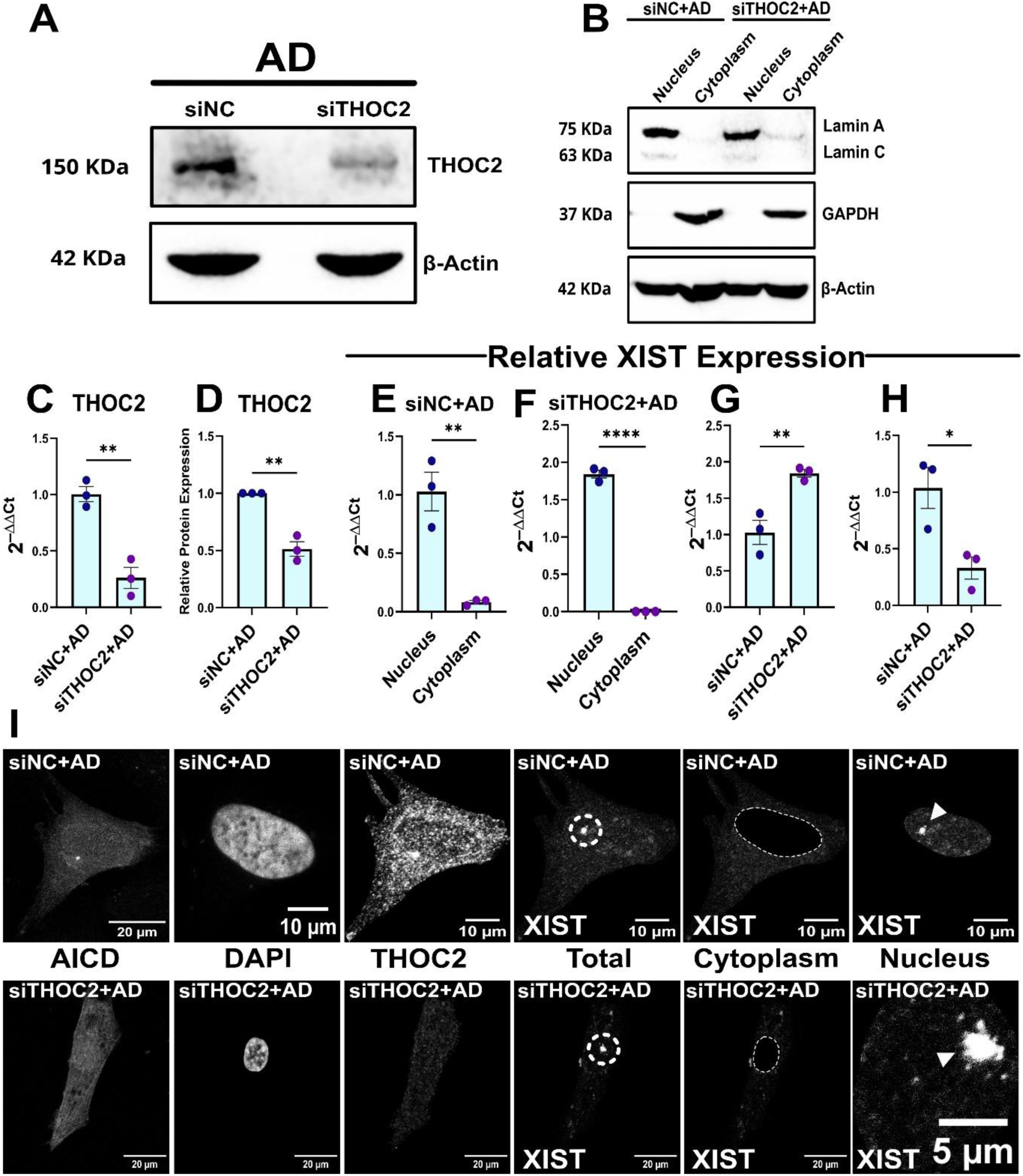
THOC2 regulates the nucleocytoplasmic distribution of XIST in AD. (A) Representative immunoblot confirming THOC2 knockdown in AICD+Aβ-treated AD cells following siTHOC2 transfection compared with the negative control (siNC). β-Actin served as the loading control. (B) Western blot validation of nuclear and cytoplasmic fraction purity using Lamin A/C as the nuclear marker and GAPDH as cytoplasmic markers. (C, D) RT-qPCR and immunoblot quantification confirming significant reduction of THOC2 mRNA and protein expression following siTHOC2 transfection in AD cells. (E, F) Subcellular fractionation followed by RT-qPCR showing the relative nuclear and cytoplasmic distribution of XIST in siNC- and siTHOC2-transfected AD cells, respectively. THOC2 knockdown resulted in increased nuclear retention and reduced cytoplasmic localization of XIST. (G) Quantification of nuclear XIST expression showing significantly increased nuclear accumulation of XIST following THOC2 depletion. (H) Quantification of cytoplasmic XIST expression demonstrating significantly decreased cytoplasmic XIST following THOC2 knockdown compared with siNC-transfected AD cells. (I) Representative RNA-FISH images showing XIST localization in siNC- and siTHOC2-transfected AD cells. THOC2 knockdown reduced cytoplasmic XIST signal and promoted nuclear accumulation, consistent with the subcellular fractionation results. Dashed outlines indicate the nuclear boundary, and arrowheads denote representative nuclear XIST accumulation. Data are presented as mean ± SEM. *p < 0.05, **p < 0.01, ****p <0.0001.

Consistent with the fractionation results, RNA-FISH analysis revealed a redistribution of XIST following THOC2 knockdown, with reduced cytoplasmic signal and increased nuclear accumulation compared to negative control treated AD cells (Figure 7I). Collectively, these results demonstrate that THOC2 promotes the cytoplasmic localization of XIST, supporting a functional role for THOC2-mediated TREX activity in XIST’s nucleocytoplasmic distribution.

### Cytoplasmic XIST helps maintain EZH2 and THOC2 in the AD nucleus via the miR186/XIST axis

The nuclear XIST was found to be upregulated in AD compared to the control (Figure 2.F). However, the EZH2-bound XIST was significantly downregulated (Figure 3.E). In the case of THOC2, XIST showed increased association in AD compared to the control (Figures 6.H). Since XIST’s functionality has traditionally been studied within nuclear territories, its cytoplasmic fate, particularly regarding miRNA sponging, remained unexplored [37]. Previous studies have shown that the miR-186/XIST sponging interaction plays a crucial role in mediating XIST-based regulation [17]. We aimed to investigate whether this axis is essential for modulating XIST’s nuclear functions involving EZH2/XIST and THOC2/XIST associations in AD.

Initially, both EZH2 and THOC2 mRNAs were found to be significantly downregulated in AD when XIST was knocked down (Figure 8.A-C). XIST-KD followed by western blots showed similar trends at the protein level as well (Figure 8.D, L, O). Thus, elevated XIST levels were found to be essential in maintaining the expression levels of these two XIST-interacting proteins. Next, we aimed to determine whether the maintenance of EZH2 and THOC2 proteins by XIST could be mediated through its interactions with miR186.

**Figure 8.**
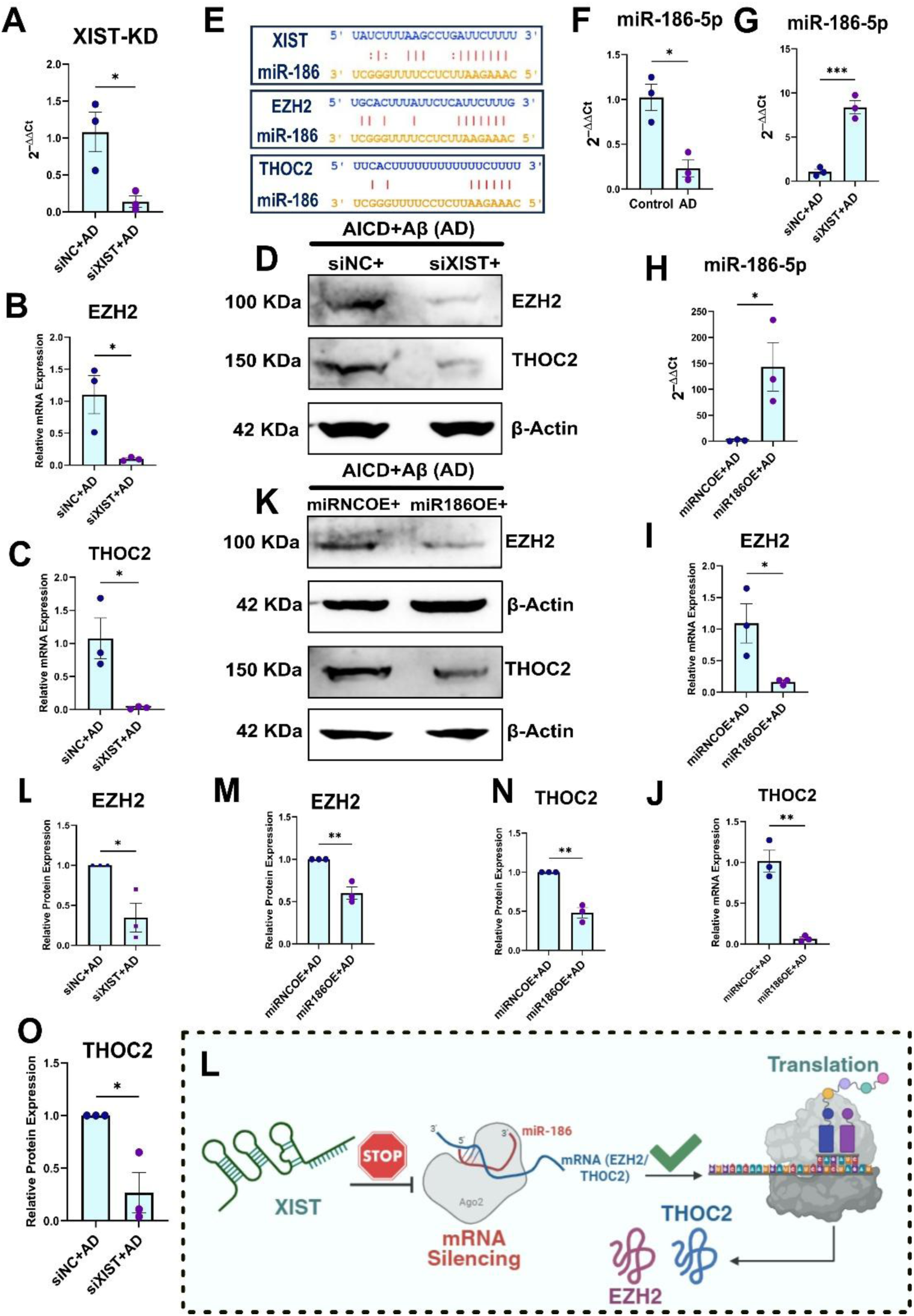
XIST regulates EZH2 and THOC2 expression through miR-186 in the AD model. (A) qRT-PCR confirming efficient knockdown of XIST in AICD+Aβ-treated AD cells following siXIST transfection. (B, C) qRT-PCR analysis demonstrating significantly reduced EZH2 and THOC2 mRNA expression, respectively, following XIST knockdown. (D) Representative immunoblots showing decreased EZH2 and THOC2 protein expression after XIST knockdown in AD cells. β-Actin served as the loading control. (E) Predicted binding sites of miR-186 on XIST, EZH2, and THOC2 transcripts obtained from StarBase. (F) qRT-PCR showing reduced miR-186-5p expression in AD cells compared with controls. (G) XIST knockdown significantly increased miR-186-5p expression in AD cells. (H) qRT-PCR confirming successful overexpression of miR-186-5p in AD cells. (I, J) qRT-PCR demonstrating significantly reduced EZH2 and THOC2 mRNA expression, respectively, following miR-186 overexpression. (K) Representative immunoblots showing decreased EZH2 and THOC2 protein expression after miR-186 overexpression in AD cells. β-Actin served as the loading control. (L, M) Quantification of EZH2 protein expression following XIST knockdown and miR-186 overexpression, respectively, demonstrating significant reductions in EZH2 protein levels. (N, O) Quantification of THOC2 protein expression following miR-186 overexpression and XIST knockdown, respectively, demonstrating significant reductions in THOC2 protein levels. (P) Schematic illustrating the proposed mechanism whereby XIST functions as a competing endogenous RNA to sequester miR-186, thereby limiting miR-186-mediated repression of EZH2 and THOC2. XIST depletion or miR-186 overexpression restores miR-186 activity, resulting in mRNA silencing and reduced EZH2 and THOC2 protein expression. Data are presented as mean ± SEM. *p < 0.05, **p < 0.01, ***p <0.001 (unpaired t-test).

The StarBase v2.0 database [38] search identified miR-186 binding sites in the mRNA transcripts of EZH2 and THOC2 (Figure 8.E). In AD, miR-186 levels were significantly reduced (Figure 8.F). Conversely, knocking down XIST in AD samples led to a significant increase in miR-186 expression (Figure 8.G), supporting the presence of the miR-186/XIST axis in AD. When miR-186 was overexpressed (Figure 8.H), there was a notable decrease in EZH2 and THOC2 mRNA (Figures 8.I, J) and protein (Figure 8.K, M, N) expression levels. This suggests that the miR-186/XIST axis regulates both EZH2 and THOC2, which then interact with XIST in the nucleus.

The mature miRNA induces post-transcriptional gene silencing via the RISC complex in the cytoplasm [39]. To further verify the novel miR186/EZH2 and miR186/THOC2 axes through cytoplasmic XIST, a biotinylated RNA pull-down experiment was performed. The miR-186 mimic with a 3’ biotin modification was transfected into the AD model. The cells were then lysed, and the miR-186-interactome was isolated from the cytoplasmic fraction using streptavidin-coated magnetic beads. The interactors were identified with qPCR (Figure 9.A). Each interaction was normalized with 10% input RNA and compared to a scrambled mimic as a negative control.

**Figure 9.**
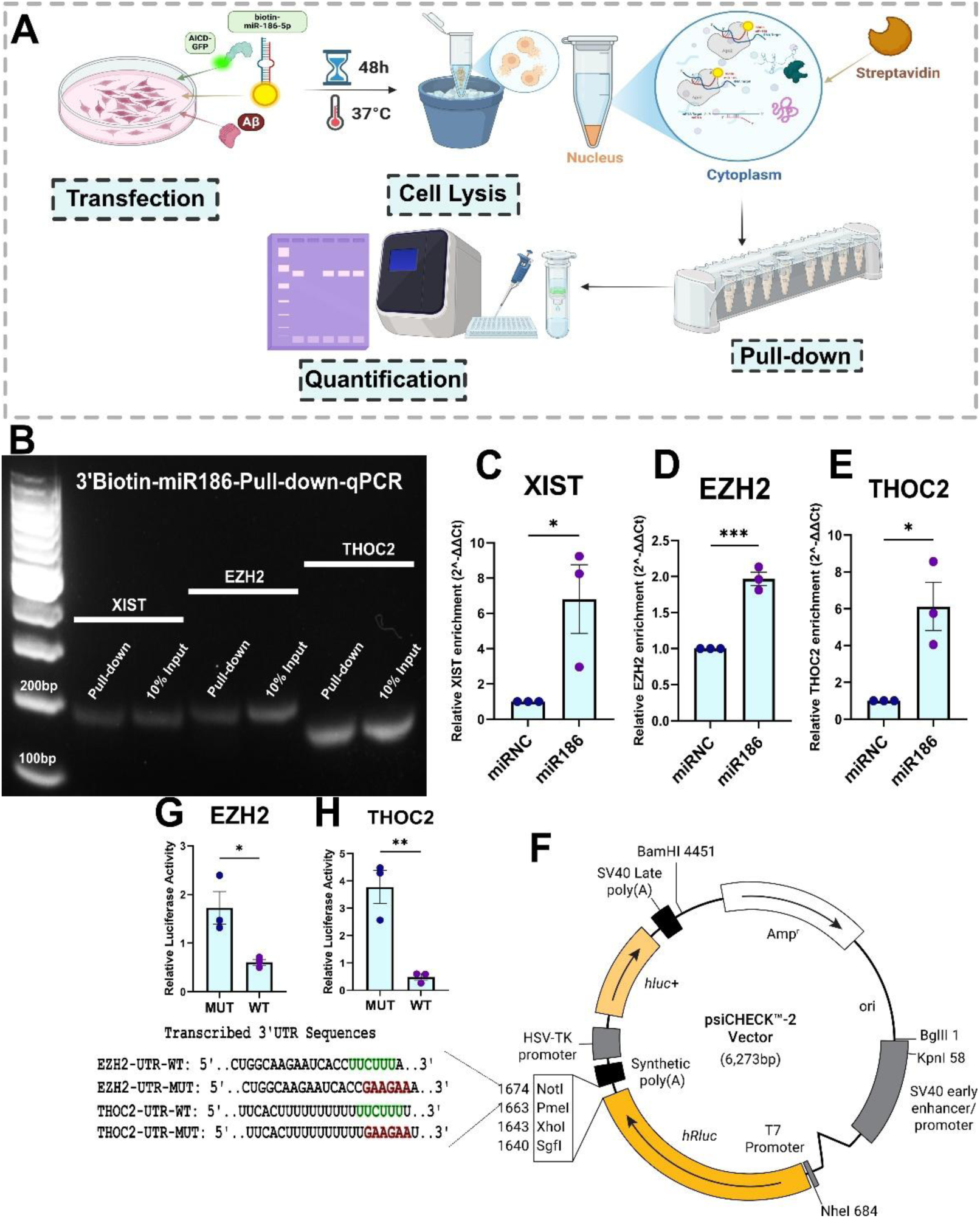
Cytoplasmic interaction of miR-186 with XIST, EZH2, and THOC2 and validation of miR-186 target binding. (A) Schematic illustration of the 3′-biotinylated miR-186 RNA pull-down assay. AICD+Aβ-treated AD cells were transfected with a 3′-biotinylated miR-186 mimic, followed by streptavidin-mediated pull-down of miR-186-associated transcripts and quantification by RT-qPCR. (B) Representative agarose gel showing successful pull-down of XIST, EZH2, and THOC2 transcripts from AICD+Aβ-treated AD cells. Ten percent input RNA served as the loading reference. (C–E) RT-qPCR quantification of pull-down products demonstrating significant enrichment of XIST, EZH2, and THOC2 transcripts in the biotinylated miR-186 pull-down compared with the miR-NC control, confirming their association with miR-186. Data were normalized to 2% input RNA. (F) Schematic representation of the psiCHECK™-2 dual-luciferase reporter vector containing the wild-type or mutant 3′UTR sequences of EZH2 and THOC2 used for validation of predicted miR-186 binding sites. (G, H) Dual-luciferase reporter assays demonstrating that miR-186 significantly reduced the luciferase activity of the wild-type EZH2 and THOC2 3′UTR reporter constructs, whereas mutation of the predicted miR-186 seed-matching sequences abolished this repression, confirming direct targeting of the EZH2 and THOC2 3′UTRs by miR-186. Data are presented as mean ± SEM. *p < 0.05, **p < 0.01, ***p <0.001 (unpaired t-test).

The pull-down experiment showed significantly strong interactions between miR-186 and XIST, EZH2, and THOC2, compared to the negative control in the AD cytoplasm (Figure 9.B-E). The cytoplasmic interactions were further verified with reduced SCARNA9 and increased tRNA-Ala expression values (Supplementary Figure S5).

Previous studies have already established the direct interaction between XIST and miR-186 using luciferase reporter assays [17]. Although bioinformatic analyses predicted conserved miR-186 binding sites within the 3′UTRs of both EZH2 and THOC2 transcripts (Figures 8.E, 9.F), these interactions have not been experimentally validated. Therefore, to determine whether EZH2 and THOC2 are direct downstream targets of miR-186, dual-luciferase reporter assays were performed using wild-type and mutant 3′UTR constructs. Co-transfection of the miR-186 mimic significantly reduced the luciferase activity of the wild-type EZH2 and THOC2 reporters, whereas mutation of the predicted miR-186 seed-matching sequences abolished this repression (Figure 9.G-H).

Together with the RNA pull-down results, these data establish that cytoplasmic XIST sequesters miR-186, thereby relieving miR-186-mediated repression of EZH2 and THOC2. Consequently, increased cytoplasmic XIST in AD sustains EZH2 and THOC2 expression, enabling the formation of the XIST-EZH2 and XIST-THOC2 regulatory axes.

## Discussion

AD continues to be a significant health issue, affecting women more than men, though the reasons remain unclear [3]. One potential contributing factor is the lncRNAs involved in regulating the disease processes [40]. These transcripts frequently exhibit inherent sex-specific biases in their expression and function [2]. The link between the lncRNA XIST and AD regulation offers new opportunities to explore the disease’s underlying mechanisms [6], especially considering its sex-specific predisposition.

XIST is well-known for its function in X-chromosome inactivation. Since female cells possess an extra X chromosome compared to males, XIST inactivates this chromosome to maintain proper gene dosage across the human genome. As a result, female cells naturally show higher levels of XIST than male cells [5], [7], [8]. It has been previously reported that XIST is markedly upregulated in AD [12], a finding supported by our snRNA-seq data and qPCR measurements from the AD cell model. This raises the question of whether XIST dysregulation may reveal female-biased aspects of AD pathophysiology.

An important limitation of the snRNA-seq analysis is the design of the GSE138852 dataset itself. Although the dataset comprises 16 biological replicates, the sequencing libraries represent pooled nuclei from pairs of donors, and the female subset contains only one pooled AD library and one pooled control library. Consequently, female-specific differential expression analyses are necessarily exploratory and cannot support donor-level pseudo-bulk analysis. Therefore, the snRNA-seq findings were interpreted as hypothesis-generating and were corroborated using independent validation studies.

The upregulated XIST from the snRNA-seq dataset (GSE138852), along with its co-expressed genes, revealed the downstream pathways impacted by this lncRNA in AD. Notably, XIST appears to promote transcriptional silencing through heterochromatin formation, while also regulating functions related to synaptic transmission and protein translation. This indicates that XIST might play an important role in AD pathology. These effects could clarify why higher XIST levels in females can be associated with an increased risk of AD.

XIST has traditionally been regarded as a nuclear-enriched RNA [41]. Recent evidence suggests the presence of cytoplasmic XIST under pro-inflammatory stress conditions [16]. Another study has previously reported the presence of cytoplasmic XIST in AD [6]. However, the functions of this cytoplasmic XIST pool remain largely unexplored. Using our cell model, we performed fractionation-qPCR and RNA-FISH to quantify cytoplasmic XIST and its contribution to overall XIST-mediated dysregulation. Our data indicate that in AD, both nuclear and cytoplasmic XIST are significantly upregulated, with a more pronounced increase in cytoplasmic XIST levels compared to nuclear XIST.

In the nucleus, the inactivated X chromosome (also known as the Barr body) is silenced through transcriptional repression. The presence of XIST across the chromosome enables the recruitment of the PRC2 complex, with the methyltransferase EZH2 as the catalytic subunit. EZH2 methylates the Lysine 27 residue of H3 histone proteins in the chromatin, mediating the transcriptional silencing of the chromosome [9], [10]. In AD, EZH2 levels were found to be significantly reduced. When XIST is pulled down through RIP with an anti-EZH2 antibody, the qPCR data suggests reduced XIST enrichment in AD compared to control. This is probably a result of reduced levels of EZH2 protein itself, which is resulting in decreased EZH2-associated XIST during RIP. This could substantially impact the heterochromatinization machinery of the Barr body. The 3D colocalization analyses reveal altered H3K27me3 marks on the inactivated X chromosome, which would, in turn, affect the X-linked genes in AD and their aberrant expression from the inactivated X chromosome in females. As epigenetic reactivation of Xi is seen in females with aging, the present study points toward the need of similar approaches to study the Xi heterochromatin and its alterations. [42], [43].

Supporting this hypothesis, our analysis of an independent female neuronal snATAC-seq dataset demonstrated a global increase in chromosome X chromatin accessibility in AD, accompanied by increased accessibility at X-linked promoter regions. These observations are consistent with a relaxation of the normally repressive chromatin state maintained by XIST and PRC2 on the Xi. However, because the snATAC-seq dataset used, do not distinguish the active and inactive X chromosomes, the extent to which the increased accessibility specifically reflects changes on the Xi cannot be directly determined.

One such X-linked gene is *THOC2*, part of the TREX complex. TREX regulates the mRNA export machinery, guiding the fate of transcripts [21]. Recent reports suggest that this machinery also facilitates the export of lncRNA from the nucleus [19], leading to the hypothesis that, with an altered epigenetic landscape, XIST could influence the abnormal upregulation of THOC2 in AD. The ChIP-qPCR studies demonstrate that the deposition of H3K27me3 on the *THOC2* promoter promotes transcriptional repression of the gene. Similar trend is followed with the THOC2 mRNA expression being downregulated in AD. However, at the protein level, THOC2 is found to be significantly upregulated in AD. This apparent discrepancy between THOC2 mRNA and protein abundance suggests that transcriptional regulation alone is insufficient to explain THOC2 expression in AD. While increased H3K27me3 deposition at the THOC2 promoter is consistent with reduced transcription, the concomitant reduction of miR-186 via XIST provides an additional post-transcriptional regulatory layer that may partially offset transcriptional repression. Nevertheless, the relative contributions of altered mRNA stability, translational efficiency, and protein turnover to THOC2 protein accumulation remain to be determined. Furthermore, ChIP-qPCR studies targeting H3K4me3 and H3K27ac could help explore how these chromatin modifiers affect the transcriptional activity of the *THOC2* promoter. The RIP-qPCR studies targeting THOC2 protein show heightened THOC2-bound XIST, suggesting a novel XIST/TREX RNA-protein complex. This upregulated THOC2-bound XIST pull-down could be due to the heightened protein expression of THOC2 in AD. Thus, for the first time, XIST can be linked with the nuclear RNA export pathways. It can further be hypothesized that this interaction may play an essential role in XIST’s own nuclear export.

To further test that hypothesis THOC2 was knocked down in AD conditions and subcellular fractionation-qPCR was performed to measure XIST levels in nucleus and cytoplasm. In knockdown conditions AD nucleus showed higher XIST levels, while the cytoplasmic XIST reduced significantly. This proves that THOC2 and TREX in general could be involved in XIST’s unusual cytoplasmic presence and nucleocytoplasmic distribution.

One of the most important pathways through which lncRNAs confer their functions is by sponging miRNAs [44]. The miRNA-186 has been previously reported to suppress BACE1, potentially suppressing the amyloidogenic pathways [18]. The expression of this miRNA is downregulated in AD. Previously, the miR186/XIST axis has been validated through luciferase assays in cancer [17]. However, how this axis would act in a neurodegenerative scenario was of significant interest. Furthermore, the role of cytoplasmic XIST in AD being unclear, it was of considerable interest to see whether the cytoplasmic XIST pool interacts with miR-186 in AD to perform its neurodegenerative functions. The miRNA pull-down studies from the cytoplasmic fraction revealed XIST’s significant association with miR-186 in the cytoplasm and the presence of two novel targets of miR-186, viz., EZH2 and THOC2. Upregulating miR-186 downregulated both these genes, while XIST knockdown upregulated miR-186 levels, resulting in the downregulation of both EZH2 and THOC2. While the nuclear H3K27me3 marks on the *THOC2* promoter drive its downregulation in AD, the significant downregulation of miR-186 by XIST in AD could partially outweigh the nuclear transcriptional repression, contributing to the overall upregulation of THOC2 protein in AD.

miRNA-mediated RNA silencing has traditionally been shown to occur in the cytoplasm [45]. The presence of XIST in the cytoplasmic compartment in AD sequesters miR-186, preventing the degradation of both EZH2 and THOC2 mRNA. Therefore, cytoplasmic XIST, through miR-186, maintains the nuclear pool of EZH2 and THOC2 proteins for nuclear XIST’s functions in AD.

A limitation of the present study is that the mechanistic experiments were performed in an AICD/Aβ-treated SH-SY5Y model, which captures several AD-associated molecular features but cannot fully recapitulate the complexity of human brain pathology. While independent human snRNA-seq and snATAC-seq datasets support the observed alterations in XIST expression and X-chromosome accessibility, future studies using human AD brain tissues and iPSC-derived neuronal cell models will be important to validate the XIST–EZH2–THOC2 regulatory axis in disease-relevant cellular contexts.

## Conclusion

Our research uncovers a dual function of XIST in AD. Inside the nucleus, it regulates epigenetic silencing, while in the cytoplasm, it affects miRNA availability, supporting the expression of key regulatory proteins. This twofold role may partly explain why AD is more common in women, as higher XIST levels could contribute to increased molecular imbalance. By identifying these linked regulatory pathways, the study deepens our understanding of sex-specific mechanisms in AD and highlights XIST as a promising target for neurodegenerative disease treatment.

## Data Availability

The dataset used in this study is publicly accessible at the NCBI Gene Expression Omnibus (https://www.ncbi.nlm.nih.gov/geo/) with the Accession ID GSE138852 and GSE174367.

## Acknowledgements

SS acknowledges fellowship support from the Saha Institute of Nuclear Physics (Government of India). Illustrations were Created with BioRender.com.

## Funding

This work was supported by the Department of Atomic Energy, Government of India (RSI 4002), and by a Core Research Grant from the Department of Science and Technology–Science and Engineering Research Board (DST-SERB), Government of India (CRG/2021/000678).

## Conflict of Interest

The authors declare no competing interests.

## Author Contributions

DM supervised the project and secured funding. SS and DM jointly conceived and designed the study. SS performed experiments, conducted data analysis, prepared figures and illustrations, and drafted the manuscript. DM reviewed and edited the manuscript.

## Supplementary Figures

**Figure S1.**
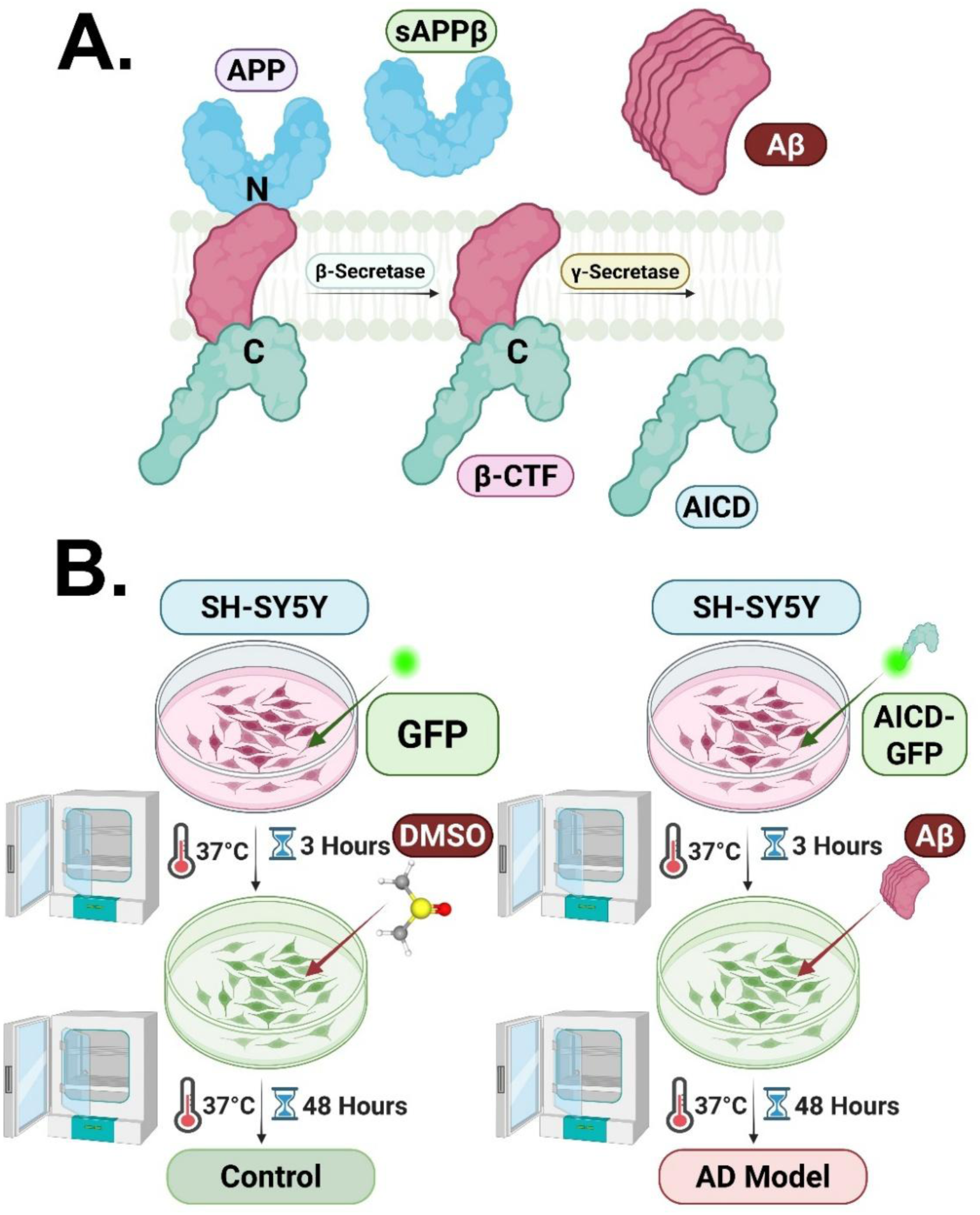
(A) In the amyloidogenic processing of APP, sequential cleavage by β-secretase and γ-secretase produces soluble APPβ (sAPPβ), the APP intracellular domain (AICD), and amyloid-β (Aβ) peptides. Aβ aggregates outside the cell, while AICD moves to the nucleus, where both fragments contribute to AD-related cellular pathology. (B) Schematic of the in vitro AD cell model. SH-SY5Y cells were transfected with either GFP (control) or AICD-GFP (AD model). After 3 hours, control cells were treated with DMSO, whereas AD cells were given Aβ peptide to mimic amyloidogenic stress. The combined presence of AICD and Aβ recapitulates the dual pathogenic signaling axis, and cells were kept at 37 °C for 48 hours before further analysis.

**Figure S2.**
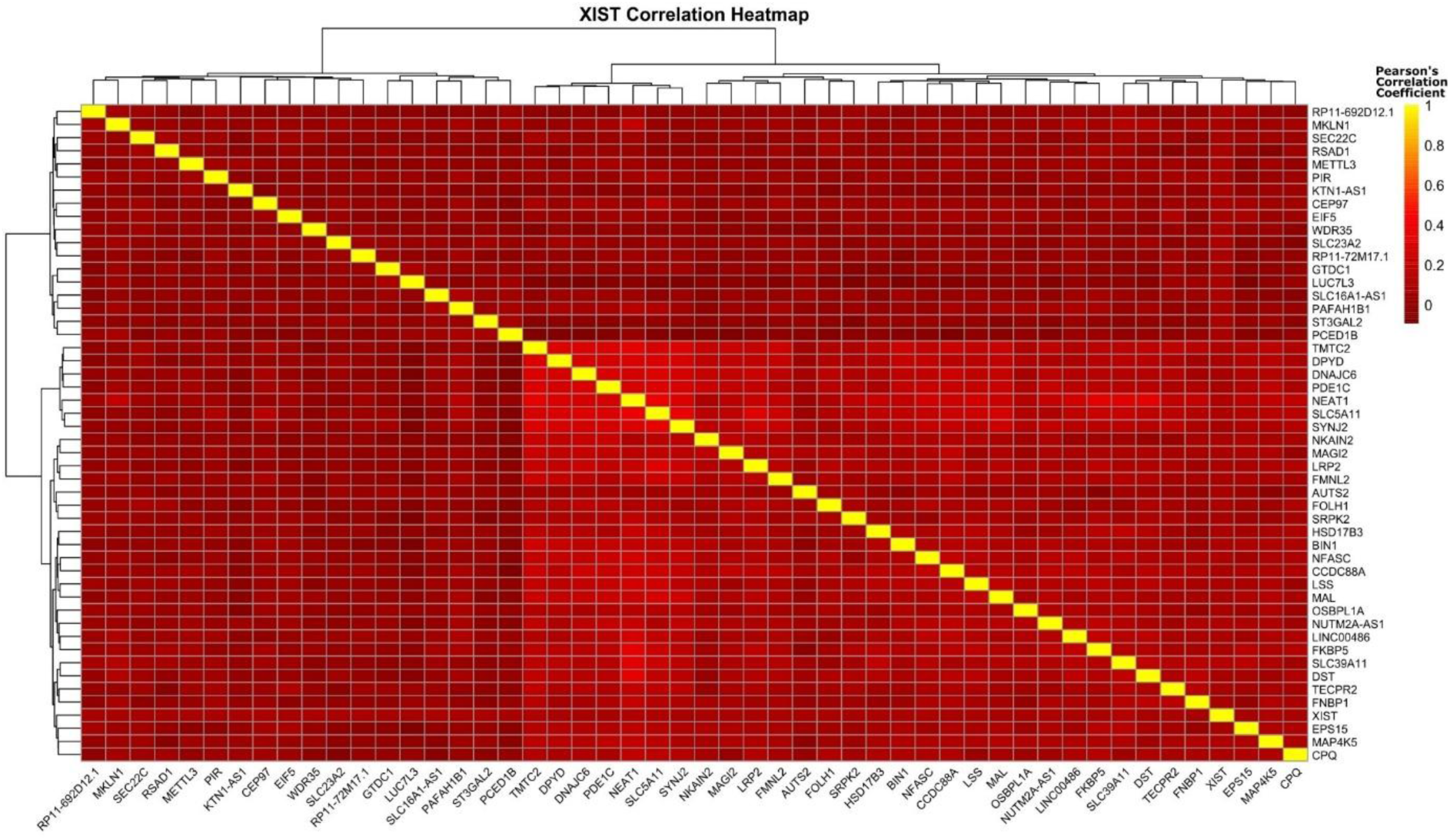
The heatmap shows the top 50 genes positively correlated with XIST in female neurons.

**Figure S3.**
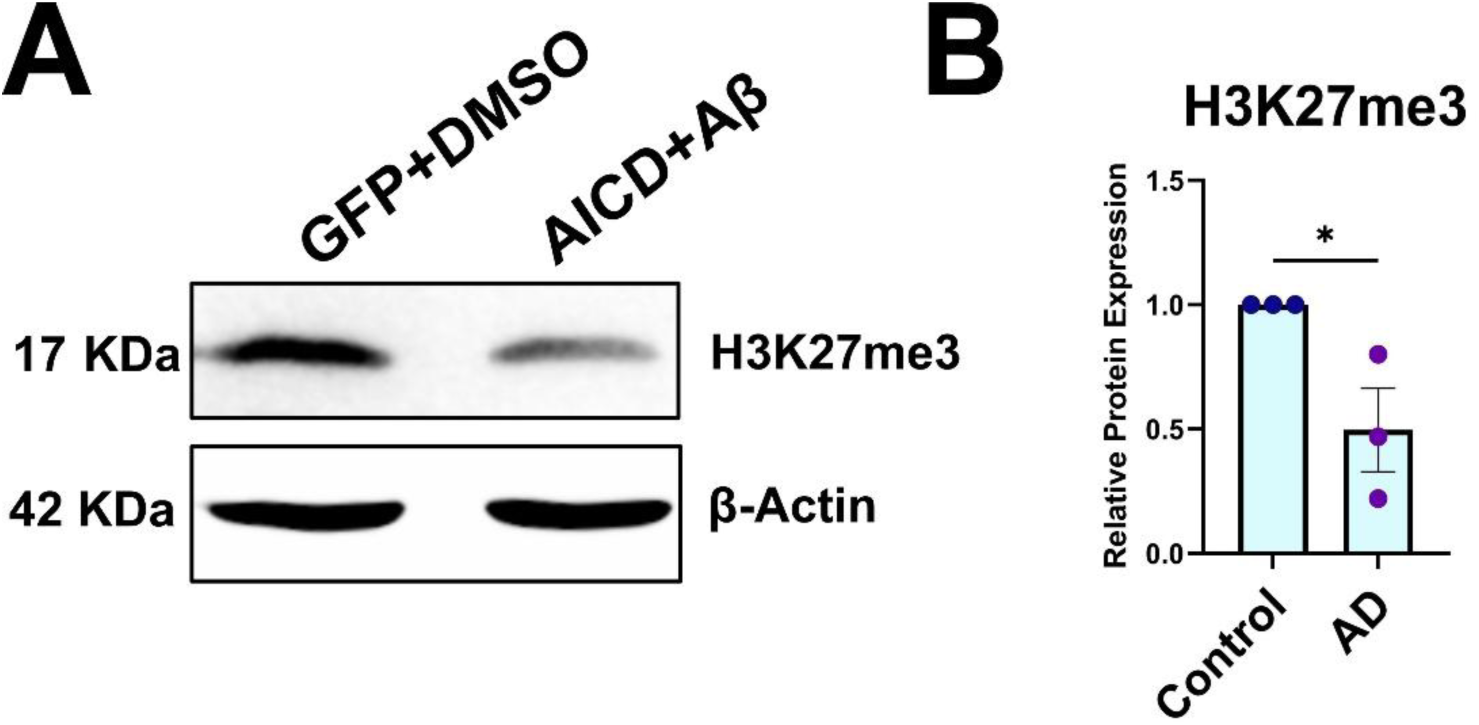
The global H3K27me3 levels were measured using western blots and was found to be significantly downregulated in AD (AICD+Aβ) compared to control (GFP+DMSO). Error bars represent mean ± SEM; significance was determined using an unpaired Student’s t test.

**Figure S4.**
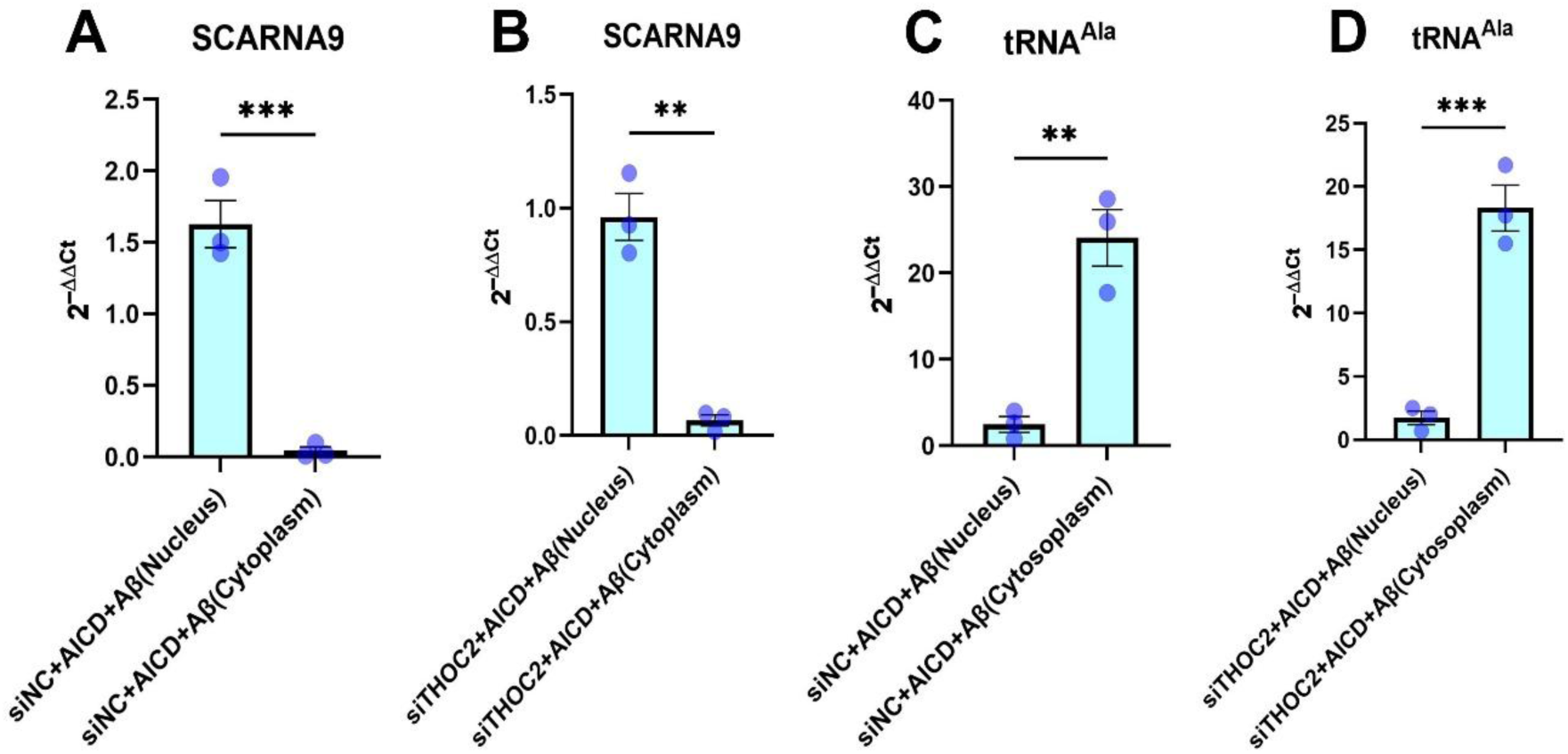
Validation of fractionation efficiency in THOC2 knock-down experiments. (A, B) SCARNA9 was significantly upregulated in the nucleus compared to the cytoplasm in both siNC and siTHOC2 treated AD cells. (C, D) tRNA-Ala was significantly upregulated in the cytoplasm compared to the nucleus. Error bars indicate mean ± SEM; significance was determined using an unpaired Student’s t test.

**Figure S5.**
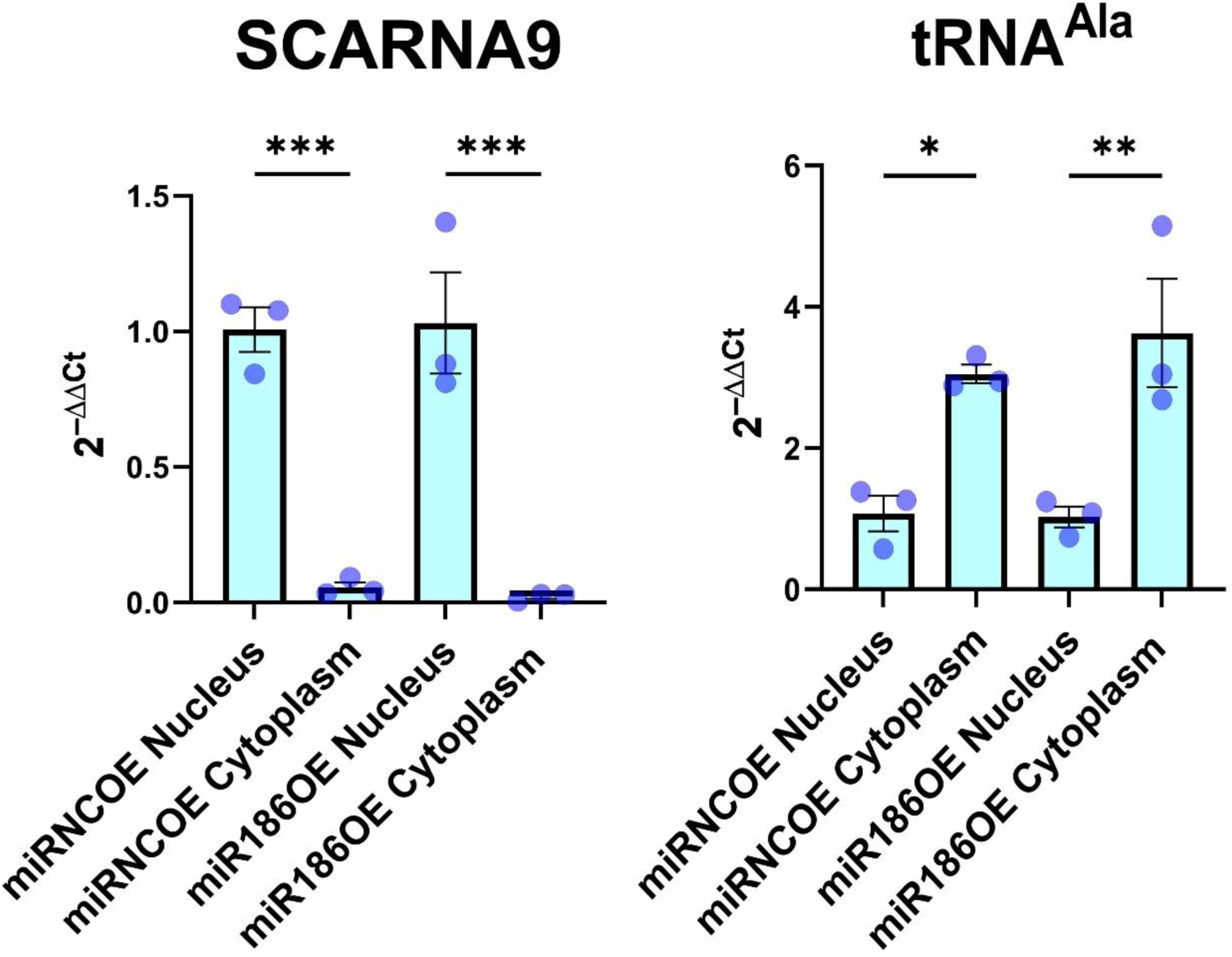
Validation of fractionation efficiency in biotinylated RNA pull-down. (A) SCARNA9 was significantly upregulated in the nucleus compared to the cytoplasm. (B) tRNA-Ala was significantly upregulated in the cytoplasm compared to the nucleus. Error bars indicate mean ± SEM; statistical significance was determined using one way ANOVA followed by Bonferroni post hoc test.

## Supplementary Table

**Table S1.**
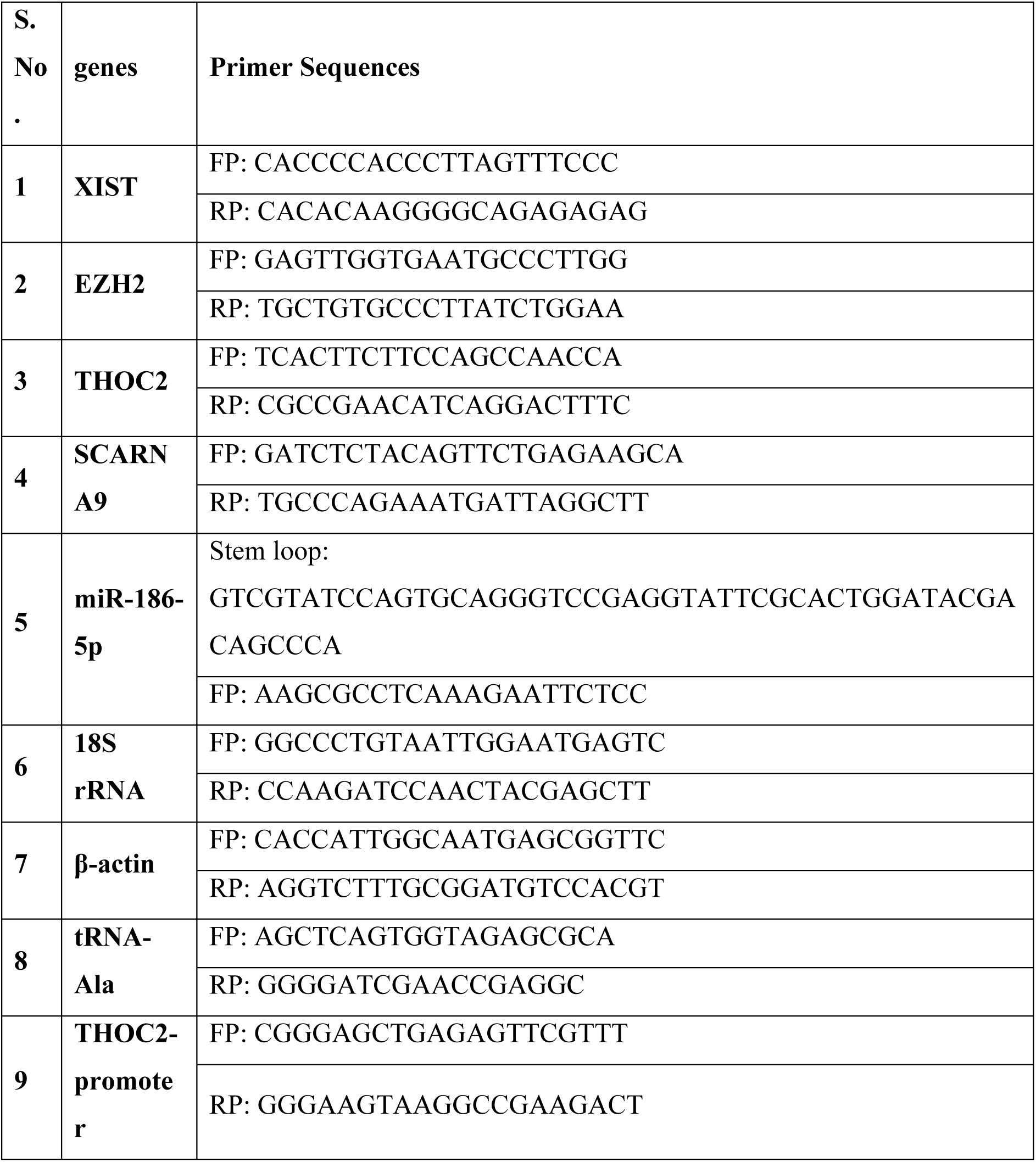
The list of primers for qPCR.

## Notes

### Competing Interest Statement

The authors have declared no competing interest.

### Summary of Updates

The updated version includes protein markers for fractionation, snATAC-seq analysis, THOC2 knockdown fractionation to validate its role in XIST's nucleocytoplasmic distribution, and dual luciferase reporter assay to further vlidate the XIST/miR-186/EZH2/THOC2 axis.

